# Tolerant crops increase growers’ yields but promote selfishness: how the epidemiology of disease resistant and tolerant varieties affect grower behaviour

**DOI:** 10.1101/2022.06.13.495875

**Authors:** Rachel E. Murray-Watson, Nik J. Cunniffe

**Affiliations:** Department of Plant Sciences, University of Cambridge, Cambridge, United Kingdom

**Keywords:** behavioural model, epidemiological model, resistance, tolerance, tomato yellow leaf curl virus

## Abstract

- Disease management often involves genetically improved crops. Resistant varieties are less susceptible, and so less likely to act as reservoirs of inoculum. Tolerant varieties can be highly susceptible, but limit yield loss for those who grow them. Population-scale effects of deploying resistant or tolerant varieties have received little consideration from epidemiologists.
- We examined how tolerant and resistant crop have opposing consequences upon the uptake of control using a behavioural model based on strategic-adaptive expectations. Growers compared last season’s profit with an estimate of what could be expected from the alternative crop type, thereby assessing whether to alter their strategy for the next season.
- Tolerant crop only benefited growers using it, decreasing yields for others. This incentivises widespread use via a negative feedback loop. Resistant crop was more widely beneficial, with reduced population-scale disease pressure leading to increased yields for all. However, this positive externality allows growers who do not deploy resistant crop to “free-ride” upon the management of others.
- This work highlights how a community of growers responds to the contrasting incentives caused by tolerant and resistant crop varieties, and how this leads to very distinct effects on yields and population-scale deployment.

## 2 Introduction

Tolerance and resistance represent the two main mechanisms underpinning the genetic control of plant disease, and the differences between the two have clear implications for epidemic management. Resistance traits are associated with a reduction in pathogen burden (Pagán & García-Arenal (2018)), and disease-resistant varieties are consequently less susceptible and/or infectious than unimproved varieties. By contrast, tolerant varieties can be infected and maintain high pathogen burdens, meaning that infected plants can transmit infection at high rates, but their yield remains largely unaffected (Pagán & García-Arenal (2018)). The two traits can be difficult to distinguish during breeding programmes, as both tolerance and resistance characteristics preserve yields when a plant is infected (Rahman et al. (2021)). Partial, or quantitative, resistance where the host does not completely restrict viral load but has a lower yield loss (Marchant et al. (2020), French et al. (2016)), is also more common than complete (quantitative) immunity (Corwin & Kliebenstein (2017)), further complicating the distinction between tolerance and resistance in breeding.

Epidemiological models have long been used to identify strategies that optimise the deployment of resistant crop (reviewed in Rimbaud et al. (2021)),Very often the focus is pathogen evolutionary dynamics and the breakdown of resistance traits (van den Bosch & Gilligan (2003), Fabre et al. (2012), Watkinson-Powell et al. (2020), Rimbaud et al. (2018)). This builds on a long history of models aiming to explain gene-for-gene polymorphisms in host and pathogen populations, stretching back to theoretical work which is now nearly fifty years old (Leonard (1977)) although is still of current interest (Tellier & Brown (2007), Clin et al. (n.d.), Hamelin et al. (2022)). However, no studies have compared the epidemiological consequences of using tolerant versus resistant crop at the population scale, despite tolerant and resistant crop having significantly distinct effects on other growers. There has also been no consideration of factors incentivising growers to deploy one or other of these possible disease controls.

though no studies to date have compared the epidemiological consequences of using tolerant vs. resistant crop. Recent models have also incorporated pathogen evolutionary dynamics and the breakdown of resistance traits. However, in our study, we only consider the epidemiological, not evolutionary, effects of the use of resistant or tolerant crop.

In economics, externalities are the effect of an action by one party on other parties (Gersovitz (2014)). Previous work has shown that in disease management schemes, prevention measures such as vaccinations generate enough *positive externalities* that they disincentivise others from partaking in control (Geoffard & Philipson (1997), Ibuka et al. (2014)). By lowering the infection burden, this type of control increases the probability that those not using the control scheme will be “successful free-riders” (that is, gain the benefit of control without paying any of the costs, Bauch & Bhattacharyya (2012)). In plant epidemiology, resistance traits lower the prevalence of infection in the system as a whole and thus have the same impact as vaccinations by decreasing the probability of infection for non-resistant crop. Previous theoretical studies have shown that not every grower need plant resistant crop for the benefits to be felt across a community of growers (van den Bosch & Gilligan (2003), Lo Iacono et al. (2013), Vyska et al. (2016)). This same result applies to other mechanisms of control. Though high participation in citrus health management areas (CHMAs; voluntary schemes established in the United States to combat citrus greening via methods such as synchronous spraying of pesticides) is correlated with better outcomes, complete participation is not needed to see improved yield outcomes (Singerman et al. (2017)).

Similarly, the majority of the benefits conferred by *Bt* -resistant maize in the United States were experienced by those that did not themselves use the improved crop and were consequently free-riding off of the actions of others (Hutchison et al. (2010)). Reducing the degree of resistance could diminish these positive externalities, but ultimately some growers planting resistant varieties still acts as a disincentive to other individuals to practice disease management.

Disease tolerance, by contrast, should have the opposite effect to resistance. Tolerant crops do not restrict pathogen replication (Pagán & García-Arenal (2018), Kause & Ødegård (2012)) and so do not reduce the probability of infection for others. Tolerant crops, however, can sustain such pathogen burdens without the same degree of yield loss as unimproved crops (Pagán & García-Arenal (2018)). Thus, growers who use tolerant crops will experience the benefit whilst generating *negative* externalities for those around them by maintaining a high infection pressure (Hozé et al. (2018)). This effect may weaken other disease management efforts, increasing the negative consequences for others (see Earn et al. (2014) for an example in human disease epidemiology where it was found that tolerance-based therapies for chronic infections increase population-level mortality as asymptomatic carriers circulate in the population). Tolerant crops may be asymptomatic and thus have a reduced probability of being visually detected and rogued (removed) in comparison to unimproved crops (Sisterson & Stenger (2018)), again allowing an increase in infection pressure. This incentivises others to use tolerant crops, too, to minimise their losses and could lead to overall higher participation in control schemes.

Game theory is an economic tool used to examine strategic decision making amongst interacting parties (Morris (2012)). As an individual’s response to the threat of disease is highly contingent on the actions of others. For example, an individual’s choice to vaccinate may depend on how many others in the population have been vaccinated (e.g. Bauch & Earn (2004), Bauch & Bhattacharyya (2012)). Game theory has been increasingly used to better understand some of the drivers of human behaviour in response to an epidemic (Chang et al. (2020)). Models derived from game-theoretic principals have also been used to examine the behaviour of growers who are making decisions about crop management (Milne et al. (2016), McQuaid et al. (2017a), Saikai et al. (2021), Murray-Watson et al. (2022)). Here, we use a game-theoretic model to examine the effects of the contrasting externalities of tolerant and resistant crop.

To examine the differing effects of disease tolerance and resistance on the profits and, consequently, the behaviour of growers, we employ *Tomato Yellow Leaf Curl Virus* (TYLCV) as a case study. Tomato (*Lycopersicon esculentum*) is a globally important crop, with over 18.5 million tonnes produced in 2020 alone (Anon (2022)). TYLCV is one of the major viruses affecting tomato production worldwide (Ramos et al. (2019)), and it has been detected from East Asia to Western Europe, the United States of America and Australia. Transmitted by the sweet potato whitefly *Bemisia tabaci* (Pan et al. (2012)), infection leads to Tomato Yellow Leaf Curl Disease (TYLCD), which causes curling of the leaves, chlorosis of young leaves, flower abortion and stunting. Combined, these symptoms can cause up to 100% yield loss. Consequently, much research has been done on developing both tolerant and resistant varieties (Vidavsky & Czosnek (1998), Dhaliwal et al. (2020)), some of which have been deployed (Lapidot et al. (1997),Riley & Srinivasan (2019)).

Despite the expansive literature, and perhaps reflecting the confusion between tolerance and resistance across crop breeding, there is considerable debate as to what degree of disease resistance has been achieved in tomato breeding. Some argue that though purportedly TYLCV-”resistant” cultivars have been developed, none have complete immunity to infection (Marchant et al. (2020) and Yan et al. (2018), who describe how what they term “resistant lines” are not actually immune to TYLCV, but instead have reduced symptoms). As there is incomplete restriction of viral replication, yield loss is less extreme than in completely susceptible genotypes. Some disease-resistant accessions have been identified in wild relatives of tomato (Yan et al. (2018)) and others have produced what appear to be genuinely disease-resistant varieties of tomato (Vidavsky & Czosnek (1998)), which will possibly lead to the deployment of fully-resistant cultivars in the future.

Additional complications such as the dependence of symptom development on nongenetic factors such as when in its life cycle the host was infected (Levy & Lapidot (2008)), or even which part of the plant was inoculated (Ber et al. (1990)) again make it harder to distinguish between resistance and tolerance. However, even the incomplete restriction of viral replication means that these cultivars can be considered partially resistant, rather than tolerant. To account for the difficulties in establishing whether a cultivar is tolerant or resistant, we consider a range of parameters pertaining to both qualities. These parameters represent a continuum between tolerance and resistance characteristics, allowing us to move smoothly from one parameterisation to another.

We use this case study to investigate the following questions: (1) How does the average profit of a group of growers change when a fixed proportion of those growers are using crop that is either tolerant or resistant to disease? (2) When growers can choose which type of crop they plant, how do the initial proportion of infectious and controlled fields affect the deployment of tolerant or resistant crop? (3) How does the use of improved crop change depending on whether it is disease-tolerant or resistant?

## 3 Description

In our model of disease spread amongst a system of tomato fields, there are two available crop varieties: unimproved crop (*U*, “uncontrolled” or “unimproved”) or an improved variety (*C*, “controlled”). The latter has some degree of tolerance and/or resistance, depending on the scaling of certain epidemiological and economic parameters. We can break down this tolerance/resistance continuum into six parameters (Table 1) which relate to how TYLCV is transmitted and the losses sustained when a field is infectious.

**Table 1:** Parameters related to resistant and tolerant varieties. The distinction between the two varieties lies in how disease is transmitted and what losses are incurred when a field is infectious.

| Parameter | Meaning | Value if resistant | Value if tolerant |
| --- | --- | --- | --- |
| $\delta_\beta$ | Relative susceptibility of improved variety | 0.5 | 1 |
| $\delta_\sigma$ | Relative infectivity of improved variety | 0.5 | 1 |
| $\delta_Y$ | Relative yield of improved variety | 1 | 1 |
| $\delta_L$ | Relative losses due to disease of improved variety | 1 | 0.1 |
| $\frac{1}{\delta_\epsilon}$ | Relative latent period for improved crop | 0.5 | 1 |
| $\delta_\nu$ | Relative probability of detection for improved crop | 1 | 0.1 |

We classify fields based on their infectious states (susceptible, *S*; latently-infected, *E*; or infectious, *I*) and by the variety with which they are planted (subscripts *U* and *C*). We do not model within-field disease spread, and assume that the two control strategies are mutually exclusive: a field cannot be planted with both unimproved and improved crop.

Susceptible fields are infected with TYLCV (transmitted by viruliferous *B. tabaci*) at rate *β* or *δ*_*β*_*β* for unimproved and improved crops, respectively. Once infected (and provided they are not first harvested), fields remain latently infected (i.e. are asymptomatic and cannot transmit disease) for an average of 1*/ϵ*, days, after which they become infectious. Fields are harvested on average every 1*/γ* days, irrespective of their control type or infection status.

We also add roguing as an control mechanism enacted by all growers irrespective of the crop variety they use. Roguing infected plants is a common practice for management of TYLCV (Ioannu (1987), Ddamulira et al. (2021), Polston & Lapidot (2007), Polston et al. (1999)). In line with the our assumption regarding symptom emergence and infectivity, only infectious crops (*I*_*i*_, *i* ∈ {*U, C*}) are rogued. Visual scouting for infection occurs at time intervals of Δ.

Tomato is a climacteric fruit, meaning it can ripen once harvested from the plant (Arah et al. (2015)). Growers, therefore, that harvest before “full maturity” call still gain marketable product (Arah et al. (2015)), though there will be a yield penalty for early harvest as immature fruit have inferior flavours and are more susceptible to damage (Tolasa et al. (2021)). As this penalty of to be less than the loss due to disease, growers who detect infection in their fields will do better by prematurely harvesting all crop to prevent disease progression (reducing losses by a factor *ϕ*_*R*_).

The degree of symptom severity will differ between unimproved and improved crop. As such, each crop type will have its own probability of detection (*v* and *δ*_*v*_*v*). We presume that when the improved crop has “tolerant” characteristics, the milder symptoms result in a low probability of detection (i.e. *δ*_*v*_ *<* 1; Table 1). However, if the improved crop has “resistance” characteristics but nevertheless becomes infected, it is detected with the same probability as the unimproved crop (i.e. *δ*_*v*_ = 1).

Unimproved and controlled fields are removed at rates *µ*_*U*_ and *µ*_*C*_ respectively which are given by (Cunniffe et al. (2014)):

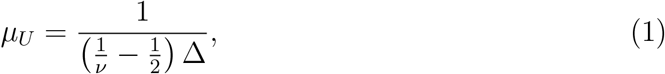

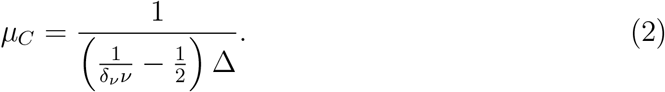

As we are modelling on the scale of a field, we presume that if a field is rogued, it is then replanted immediately with healthy crop. The epidemiological model is given by:

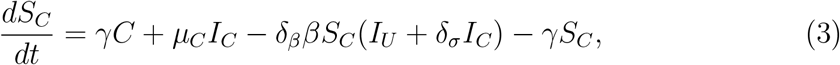

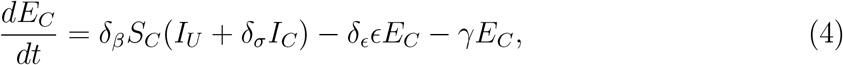

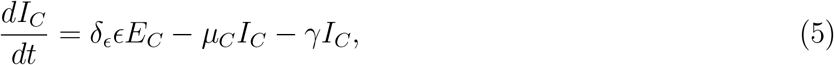

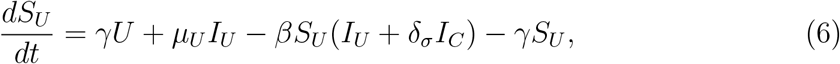

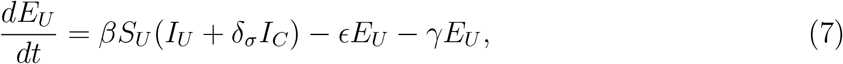

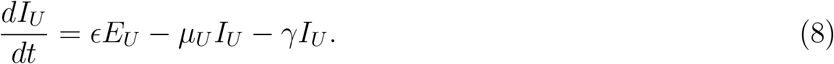

where *C* = *S*_*C*_ + *E*_*C*_ + *I*_*C*_ and *U* = *S*_*U*_ + *E*_*U*_ + *I*_*U*_.

The rate of roguing applies to the field level, but when we calculate the profits for each strategy we do so for individual growers. Roguing aims to minimise yield loss; the losses due to disease (*L*) are thus reduced by a factor *ϕ*_*R*_ *<* 1 that represents the relative benefit of harvesting an infected field before the end of the season and thus avoiding the maximum yield loss. The benefit of roguing is only accounted for in profit of the infected controllers as we assume that it is incurred immediately (i.e. roguing and replanting a rogued field occur in the same step, and all fields that are rogued are replanted).

Parameters for this model are summarised in Table 2 and initial conditions outlined in Table 3.

**Table 2:**
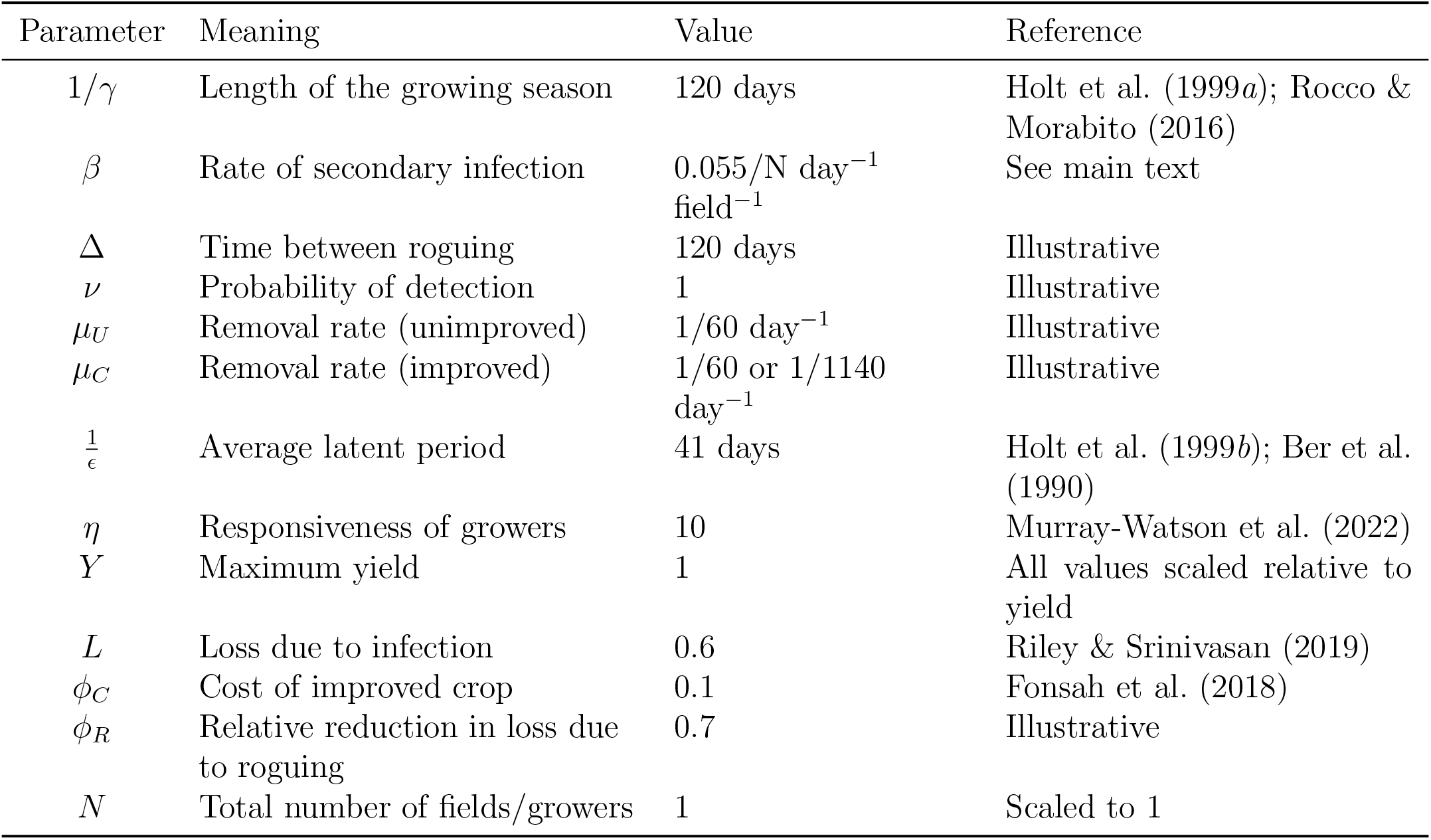
Summary of parameter values. When the parameter relates to improved crop, the first value is for tolerant crop and the second for resistant.

| Parameter | Meaning | Value | Reference |
| --- | --- | --- | --- |
| $1/\gamma$ | Length of the growing season | 120 days | Holt et al. (1999a); Rocco & Morabito (2016) |
| $\beta$ | Rate of secondary infection | $0.055/N \text{ day}^{-1} \text{ field}^{-1}$ | See main text |
| $\Delta$ | Time between roguing | 120 days | Illustrative |
| $\nu$ | Probability of detection | 1 | Illustrative |
| $\mu_U$ | Removal rate (unimproved) | $1/60 \text{ day}^{-1}$ | Illustrative |
| $\mu_C$ | Removal rate (improved) | $1/60$ or $1/1140 \text{ day}^{-1}$ | Illustrative |
| $\frac{1}{\epsilon}$ | Average latent period | 41 days | Holt et al. (1999b); Ber et al. (1990) |
| $\eta$ | Responsiveness of growers | 10 | Murray-Watson et al. (2022) |
| $Y$ | Maximum yield | 1 | All values scaled relative to yield |
| $L$ | Loss due to infection | 0.6 | Riley & Srinivasan (2019) |
| $\phi_C$ | Cost of improved crop | 0.1 | Fonsah et al. (2018) |
| $\phi_R$ | Relative reduction in loss due to roguing | 0.7 | Illustrative |
| $N$ | Total number of fields/growers | 1 | Scaled to 1 |

**Table 3:** Default initial conditions. These are scaled to be proportions (i.e. *N* = 1).

| Variable | Meaning | Value |
| --- | --- | --- |
| $S_C(0)$ | Initial proportion of susceptible controlling fields | $0.1N$ |
| $E_C(0)$ | Initial proportion of latently infected controlling fields | $0N$ |
| $I_C(0)$ | Initial proportion of infectious controlling fields | $0N$ |
| $S_U(0)$ | Initial proportion of susceptible non-controllers | $0.89N$ |
| $E_U(0)$ | Initial proportion of latently infected non-controlling fields | $0N$ |
| $I_U(0)$ | Initial proportion of infectious non-controllers | $0.01N$ |

### 3.1 Parameterisation

Where possible, parameter values were taken from the literature. A summary of the parameterisation is given in Table 2, with full details discussed in Appendix 2.

### 3.2 Growers’ profits

To determine the benefits provided by each strategy, we estimate the expected profits of a grower using a particular strategy. These are then compared with the grower’s profit from the previous season to determine if the grower should consider switching strategy. These profits account for the costs and losses associated with each crop type and infection outcome and are given by:

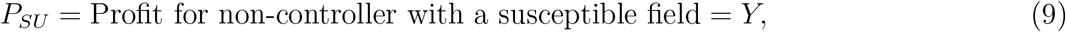

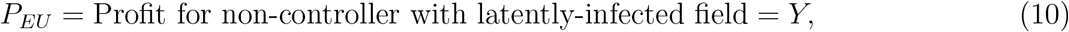

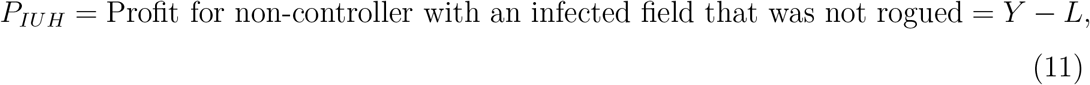

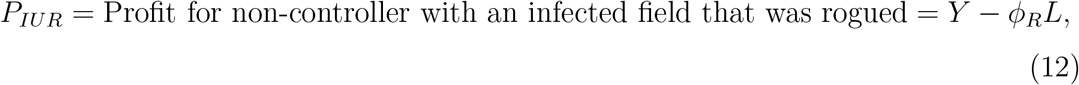

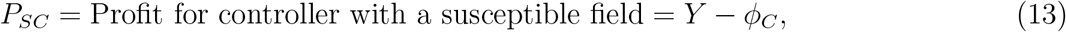

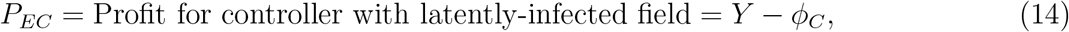

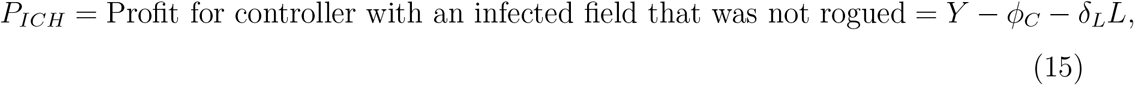

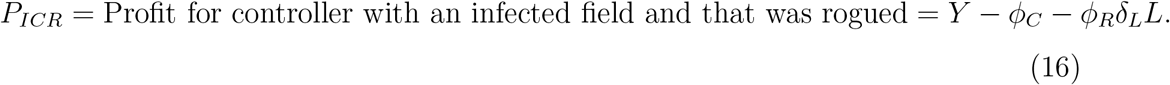

The profit for an uninfected or latently-infected field using unimproved crop (*P*_*SU*_ or *P*_*EU*_) will always the maximum achievable profit, as at the time of harvest these growers have avoided paying the cost of control or incurring any losses due to infection. We differentiate between profits of growers with infectious fields by whether or not their field was rogued before harvesting, and thus had a lower yield loss (*ϕ*_*R*_*L* or *ϕ*_*R*_*δ*_*L*_*L*).

As we only consider the case where the costs of control are less than the losses due to disease in unimproved crop (*ϕ*_*C*_ *< L*), the relative sizes of the remaining profits depend on the tolerance/resistance characteristics of the improved crop. For tolerant crop, where *δ*_*L*_*L < L*, the order of profits is:

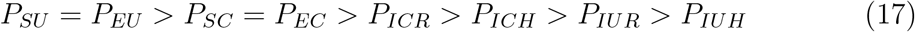

and for a resistant background is, where *δ*_*L*_*L* = *L*:

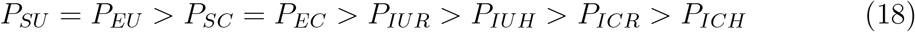

In the case where there is a fixed proportion of growers using each strategy, we can quantify the benefit each strategy provides by calculating the average profits for a grower using each crop type. To do this, we must first define the probability that an infectious field has not been rogued (*q*_*IUH*_ and *q*_*ICH*_ for unimproved and improved fields respectively):

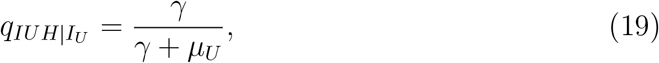

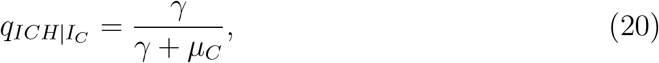

and the field has been rogued is:

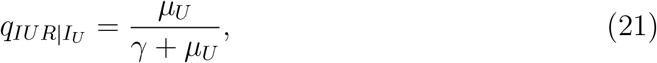

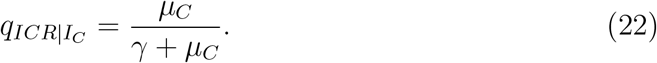

The average profits for unimproved (*P*_*U*_) and improved (*P*_*C*_) crop are given by:

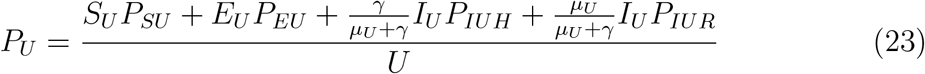

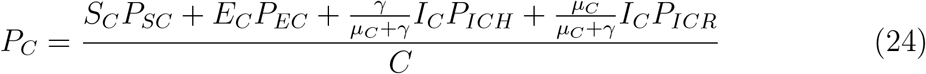

where *U* = *S*_*U*_ + *E*_*U*_ + *I*_*U*_ and *C* = *S*_*C*_ + *E*_*C*_ + *I*_*C*_.

### 3.3 Calculating expected profits

In the above, growers were assigned a control strategy at the beginning of the epidemic and could not change to the alternative, irrespective of profitability or grower preference. However, it is expected that growers will instead choose whether to control based on the perceived profitability of control, which will depend on parameters such as the cost of control and also the current risk of infection.

We use the “grower vs. alternative” mechanism for decision making, as set out in Murray-Watson et al. (2022), McQuaid et al. (2017a), Milne et al. (2016), Saikai et al. (2021). In this behavioural model, growers compare their outcome from the previous season with the expected profit of the alternative strategy (i.e. the strategy that they did not previously adopt), which in turn is based on the instantaneous probability of infection. The probability that they change strategy is then based on the magnitude of the differences between these their previous profit and the profit of the alternative strategy.

The probability of horizontal transmission for a grower using unimproved crop (*q*_*U*_) is given by:

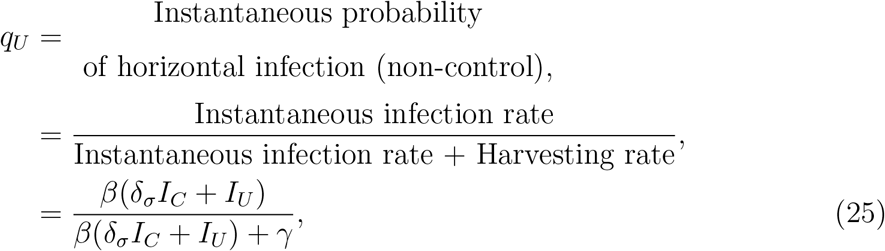

and for a grower using improved crop (*q*_*C*_) is:

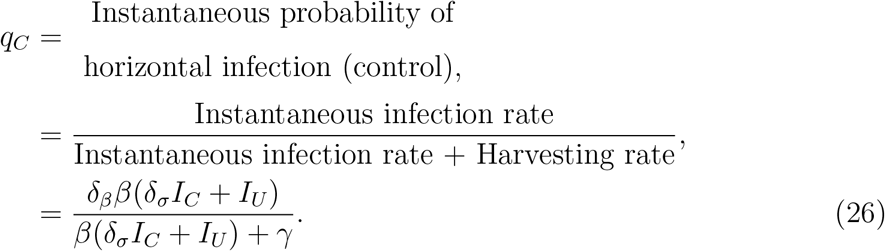

We must also consider the probability that, once infected, a grower will be latently infected (*E*) or infectious (*I*). The probabilities that a field planted with unimproved or improved crop will be latently infected at the time of harvest (*q*_*EU*_ and *q*_*EC*_) are given by:

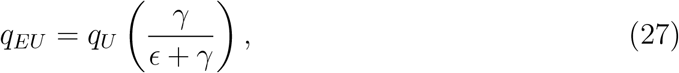

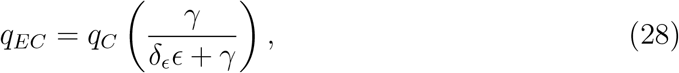

whilst the probabilities the field is infectious (*q*_*IU*_ and *q*_*IC*_) are:

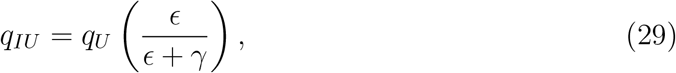

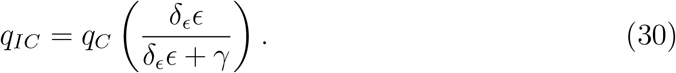

Finally, we must consider the probability that a field is infectious and then rogued (*q*_*IUR*_ and *q*_*ICR*_) before it is harvested:

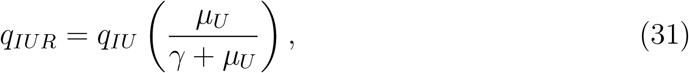

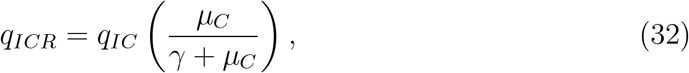

or that it is harvested (*q*_*IUH*_ and *q*_*ICH*_) before being rogued:

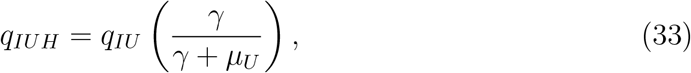

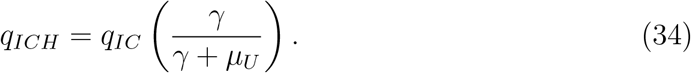

The expected profits for a non-controller, *P*_*U*_ is therefore given by:

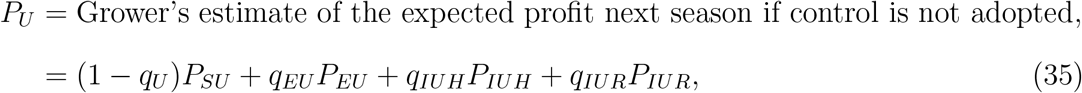

and for a controller (*P*_*C*_) it is:

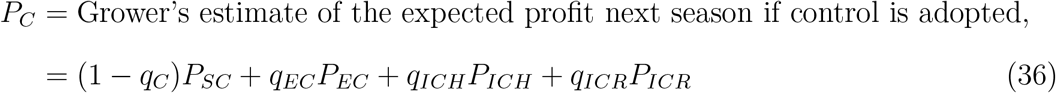

Using Equations 9 - 16, Equations 35 and 36 be further simplified:

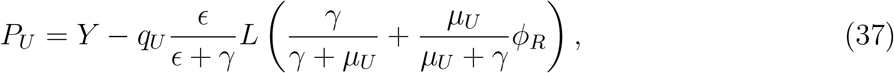

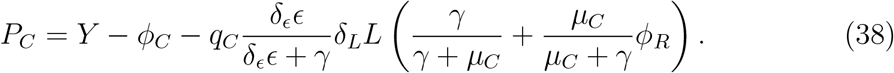

### 3.4 Switching terms based on the expected profits

From Equations 37 - 38, we can base determine the probability of a grower of outcome *i* ∈ {*S*_*U*_, *E*_*U*_, *I*_*U*_, *S*_*C*_, *E*_*C*_, *I*_*C*_} switching into the alternative strategy (Murray-Watson et al. (2022)). These “switching terms” compare the difference between the grower’s profit from the previous season with the expected profit of the strategy that they did not adopt. These differences are multiplied by a “responsiveness” parameter, *η*, which accounts for the responsiveness of growers to differences in profit (Murray-Watson et al. (2022)). If the expected profit is less than the grower’s current profit, the grower should not switch strategy. The payoff for a non-controller that harvests susceptible crop (*P*_*SU*_ and, for the default parameterisation, *P*_*EU*_ from latently-infected fields) should always be the highest as they do not pay the cost of control or losses due to disease. These growers should therefore never switch strategy. Which is the lowest payoff will depend on whether the improved crop is tolerant or resistant (Equations 17-18). If it is tolerant, the loss for tolerant crop will be less than that of unimproved crop (*δ*_*L*_*L < L*). Consequently, *P*_*IUR*_ is the lowest payoff. Conversely, if the improved crop is resistant, *δ*_*L*_*L* = *L* and the cost of control means that *P*_*ICR*_ is the lowest payoff.

The switching terms are given by:

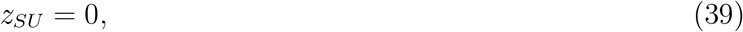

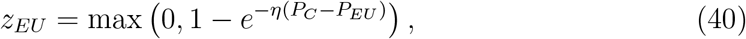

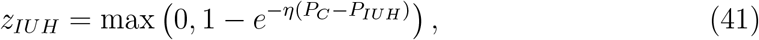

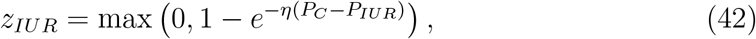

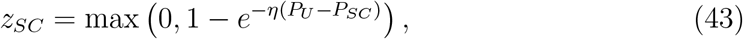

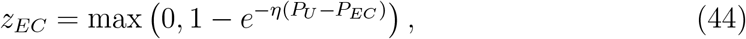

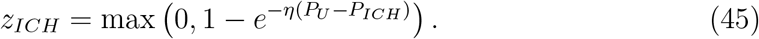

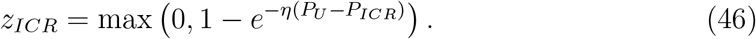

Incorporating these into the epidemiological model, we have:

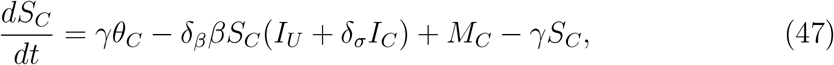

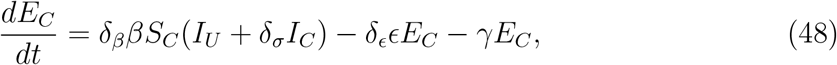

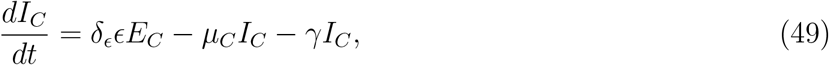

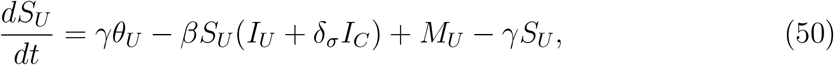

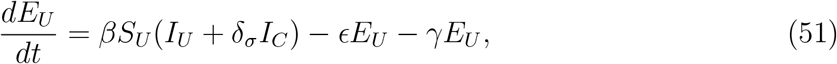

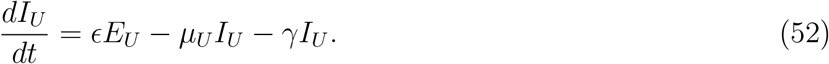

where:

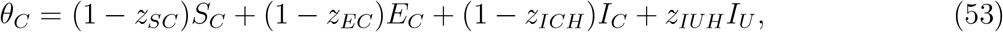

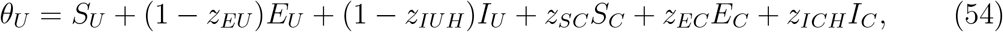

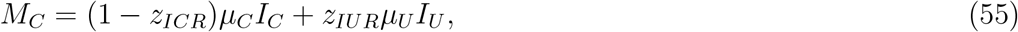

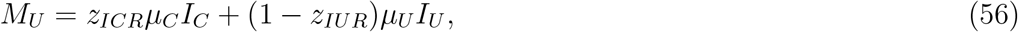

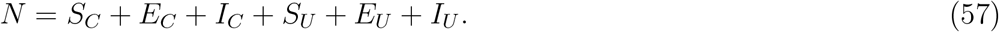

We highlight here that in Equation 54, *S*_*U*_ and *E*_*U*_ are not associated with any switching terms as for our parameterisation, the values of *z*_*S*_*U* and *z*_*E*_*U* are zero. A schematic of the model is provided in Fig. 1.

**Figure 1:**
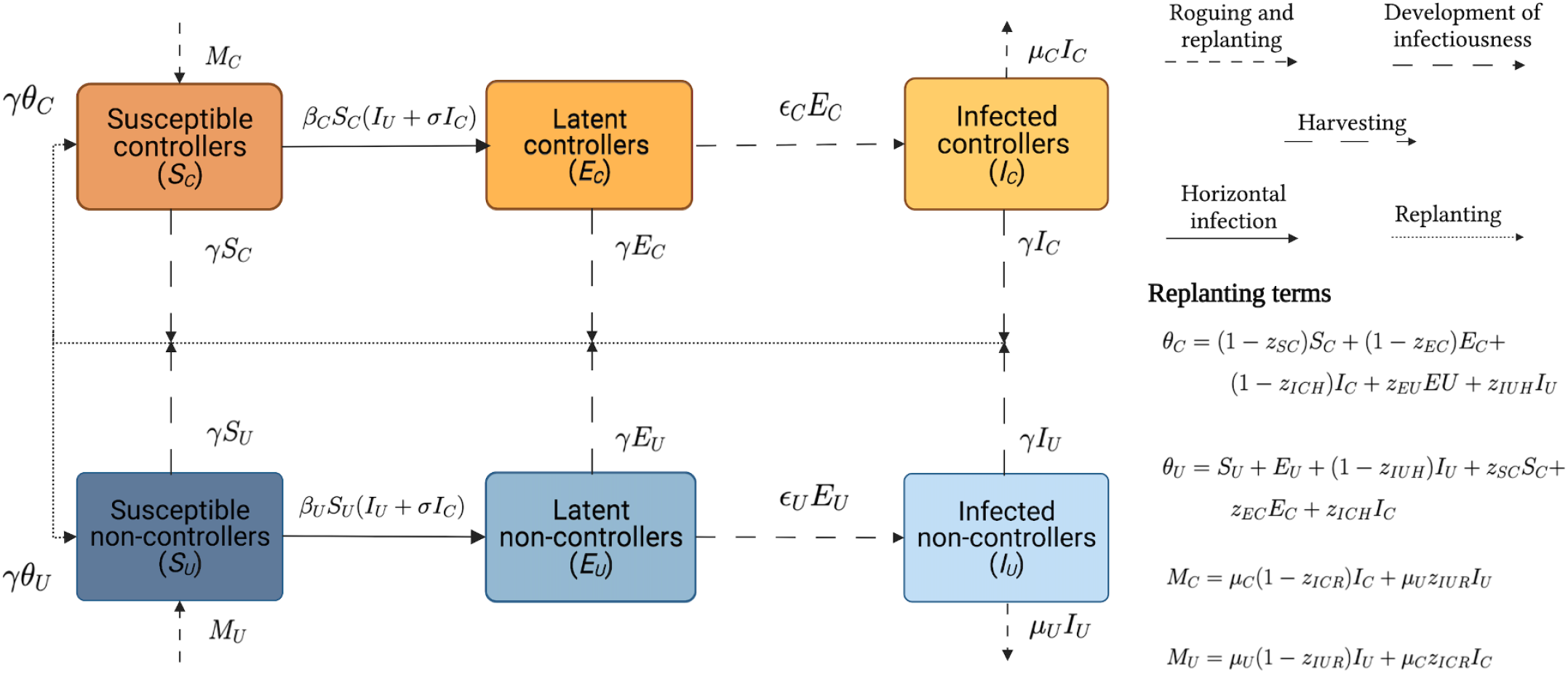
Schematic showing the structure of the model when growers can choose their strategy based on expected profits. We have two classes of grower, those who use unimproved seed (*U*) and those who use improved seed (*C*). This improved seed comes in one of two varieties: tolerant or resistant. The terms *θ*_*C*_ and *θ*_*U*_ are the rates of replanting for harvested improved and unimproved fields, whilst *M*_*C*_ and *M*_*U*_ are rates of replanting for rogued fields (Equations 53 - 56, with *M*_*C*_ + *M*_*U*_ = *µ*_*C*_*I*_*C*_ + *µ*_*U*_ *I*_*U*_). Created with BioRender.com

Which switching terms are positive or fixed at zero depends on the parameter values and the epidemiological state of the system. Due to the ordering of the payoffs, only certain combinations of positive switching terms are possible (Appendix 2).

## 4 Results

### 4.1 Q1: Externalities of tolerant and resistant crop with fixed proportions of growers

#### 4.1.1 Basic reproduction number, *R*_0_

When there is only one type of crop (improved, which can be either resistant or tolerant, or unimproved), the basic reproduction numbers are as follows:

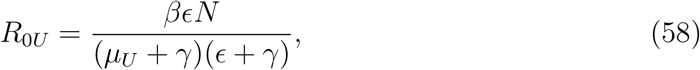

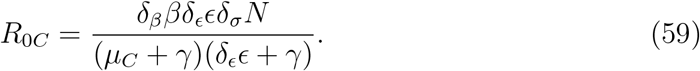

where for *R*_0*U*_, *N* = *U* and for *R*_0*C*_ *N* = *C*. In these expressions 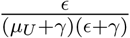 and 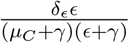 are the probabilities that a field will become infectious before it is harvested for unimproved and controlled fields respectively. The mean time spent in the *I*_*i*_ compartment is 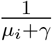 (with *i* ∈ {*U, C*}) is the mean time in the *I*_*i*_ compartment and the number of infections given caused by these infectious fields is *βU* or *δ*_*β*_*βC* for unimproved or improved respectively. The value of *R*_0*C*_ also accounts for the reduced infectivity of infectious improved crop (*δ*_*σ*_).

We use the next generation matrix method (NGM; van den Driessche (2017)) to evaluate *R*_0_ when both crop types are present at the disease-free equilibrium (i.e. (*S*_*U*_, *E*_*U*_, *I*_*U*_, *S*_*C*_, *E*_*C*_, *I*_*C*_) = (*U*, 0, 0, *C*, 0, 0)) (see Appendix 3).

*R*_0_ when both crop types are present is given by:

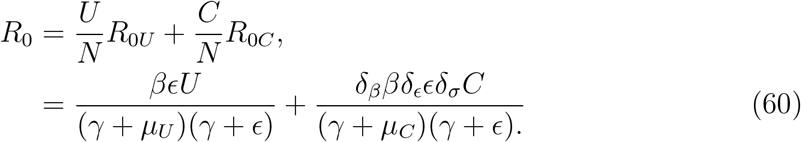

This is a combination of Equations 58 and 59, scaled in proportion with the proportion using each crop type. This is concurrent with Bosch et al. (2008) showing a linear relationship between the proportion of each crop in a mixed-cultivar system and the basic reproduction numbers for those cultivars if they were deployed in a monoculture.

#### 4.1.2 Effect of changing proportion of improved crop

We first consider the effect of an increased proportion of improved crop on the expected profits of growers of each type. Using the default parameters outlined in Table 2, we can see that an increase in the proportion of growers using tolerant crops has little impact on the expected profit of controllers (decreasing profits by ∼1%), though reduces those of non-controllers (Fig. 2a). As tolerant crops have a lower probability of being detected (*δ*_*v*_*v* = 0.1) and thus a lower removal rate (*µ*_*U*_), having more tolerant crop increases the disease pressure fewer infectious fields are removed via roguing (Fig. 2c).

**Figure 2:**
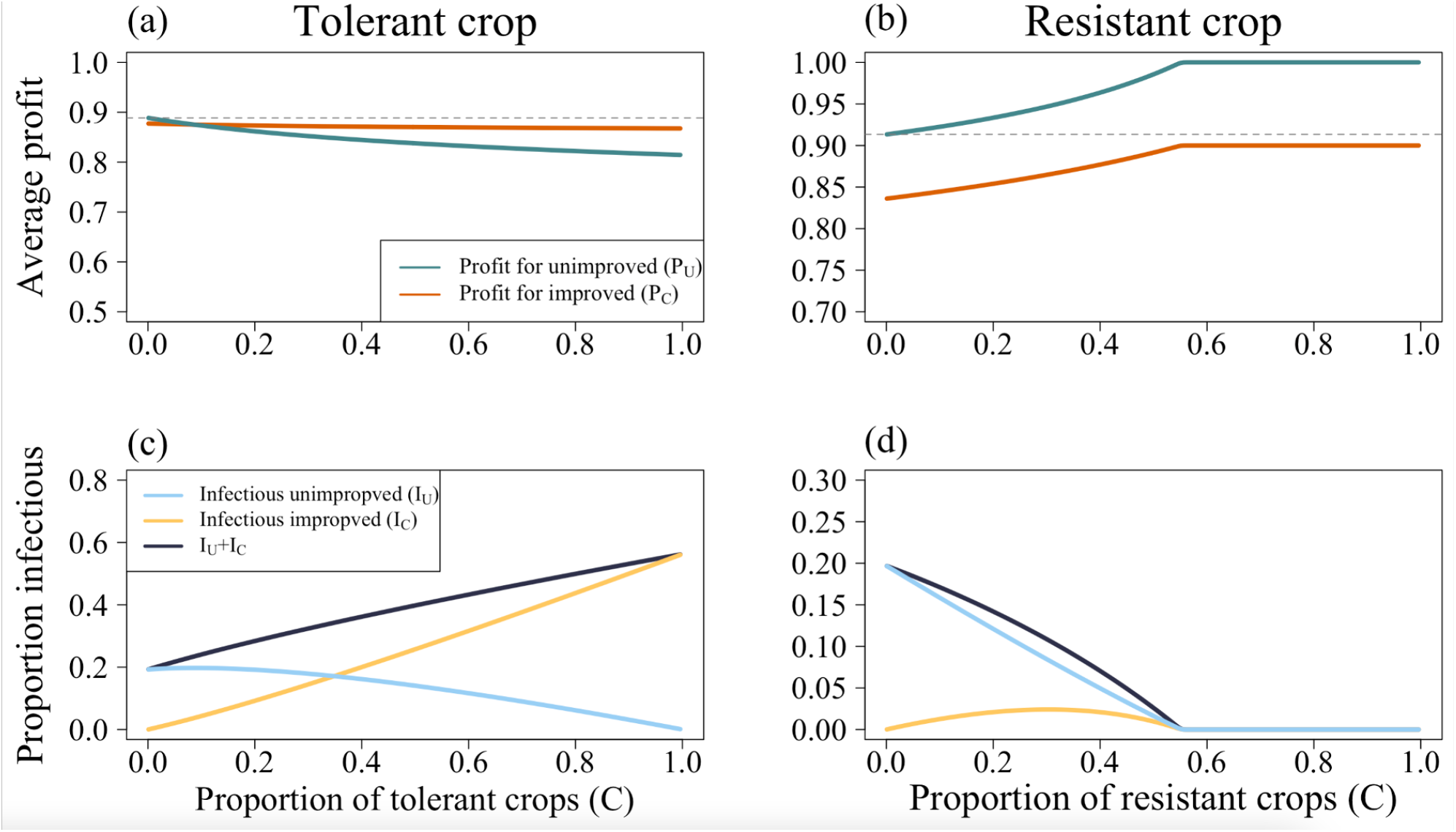
Effect of increasing proportions of improved crop on the average profit of growers using unimproved and improved crop. (a) and (c) use the tolerant parameterisation whilst (b) and (d) show the same for resistant crops. The two types of improved crop had opposite effects; as the proportion of improved, tolerant crop increased, so too did the amount of infection. This caused a decrease in profits for non-controllers. Conversely, as *C* increased when the crop was resistant, there was a decrease in the amount of infection and concomitant increases in profits. In both cases, the grey dashed line shows the average profit of a non-controller when there are no controllers (*C* = 0), which can be used to measure the externalities generated by each type of improved crop. Parameters and initial conditions are as in Tables 2 and 3 respectively.

Conversely, an increase in the proportion of resistant crops provided much greater benefits to non-controllers than to controllers (Fig. 2b). Controllers already had a relatively low probability of infection, so their average profit is already close to the maximum possible (*P*_*SC*_ = 0.85). Thus, a decrease in the probability of infection due to an increased proportion of growers using resistant crop provides little additional benefit. With a sufficient proportion of resistant crops, disease is eliminated from the system (Fig. 2d). However, in this scenario controllers had to continue paying the cost of control (*ϕ*_*C*_) even though there is little need for control, so the controllers earn less than the non-controllers.

In both of these graphs, the deviation from the average profits where there is no improved crop in the system (*C* = 0, indicated by the grey dashed line) can be seen as the magnitude of the externalities generated by each crop type. As an increase in tolerant crop causes *P*_*U*_ to decrease, it generates negative externalities. Conversely, the resistant crop reduces the probability of infection and thus generates positive externalities, increasing *P*_*U*_. The increase in *P*_*U*_ at higher values of *C* (from *P*_*U*_ = 0.91 when *C* = 0 to *P*_*U*_ = 1 when *C* = 0.99) is greater than the corresponding increase in *P*_*C*_ (from from *P*_*C*_ = 0.836 when *C* = 0 to *P*_*C*_ = 0.9 when *C* = 0.99), indicating that a greater benefit is felt by non-controllers than by controllers.

We have shown the above for only a single set of parameters, but the broad patterns are recapitulated for parameters controlling the effectiveness of the tolerant/resistant crop (namely the probability of detection of improved crop (*δ*_*v*_*v*) and the relative susceptibility of improved crop (*δ*_*β*_) (Appendix 4 Figs. 1 and 2)).

### 4.2 Q2: Effect of initial conditions on long-term outcomes in the behavioural model

The complexity of the model (in particular, the presence of the switching terms) means that explicit expressions cannot be found for the values of state variables at equilibrium. However, their values and the stability of the equilibria can be evaluated numerically.

Using the NGM method (see Appendix 5), we found the basic reproduction number for the behavioural model to be:

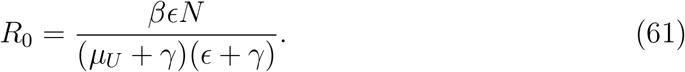

Broadly, this determines the stability of the disease-free equilibrium, as disease can only invade the system once *R*_0_ *>* 1. In some cases, however, bistability is possible and there may be multiple possible equilibria below this threshold depending on the initial conditions (Gumel (2012)).

The long-term outcomes of the model can be divided into one of four types:

- **Disease-free equilibrium:** as *R*_0_ *<* 1, there are no infected fields. As there is no risk of infection, no growers use improved crop (and therefore avoid the cost of control, *ϕ*_*C*_).
- **“No control” equilibrium:** disease is endemic, but no growers use improved crop.
- **“All control” equilibrium:** disease is endemic and no growers use unimproved crop.
- **Two-strategy equilibrium:** disease is endemic and crops of both varieties are used.

For a given parameter set, it may be that two of these equilibria are locally stable, depending on the initial conditions.

The parameterisation of the model, both in terms of whether the improved crop is tolerant or resistant and its degree of tolerance and resistance, determines the subset of possible long-term outcomes. When the improved crop was resistant, an “all control” equilibrium was not possible (Fig. 3a). The positive externalities generated by the presence of resistant crop disincentivises non-controllers from using improved crop, as they have a lowered probability of infection without themselves being infected.

**Figure 3:**
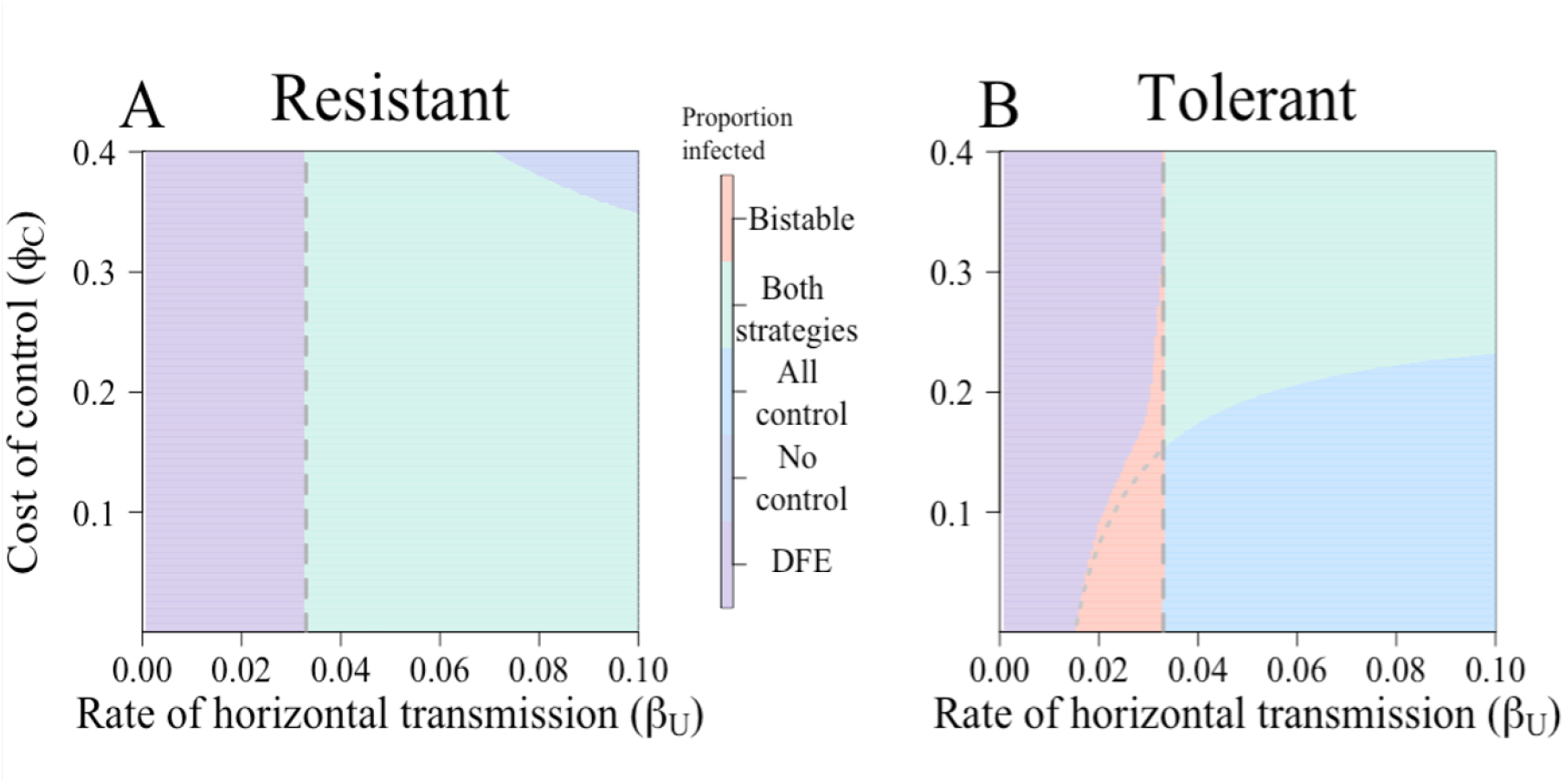
Nature of equilibria attained when improved crop is either tolerant or resistant. There are five possibilities: a “disease-free equilibrium” (DFE), with only unimproved crop; a disease-endemic, “no control” equilibrium; a disease-endemic, “all control” equilibrium; and an equilibrium where both strategies and disease are present and finally a bistable region of parameter space, where two stable equilibria are possible and so the long-term behaviour of the system depends on the initial conditions (see also Fig. 4). For both parameterisations *R*_0_ = 1 is indicated by the grey vertical dashed line at *β* = 0.0333 day ^*−*1^. (a) Equilibria for the resistant crop. Here, only three of the possible equilibria exist: at no point can there be an “all control” equilibrium. Additionally, there is no bistable region. (b) Equilibria for the tolerant crop. Now, at lower costs of control and medium-to-high values of *β*, an equilibrium where all growers use improved crop is possible. There is a bistable region for 0.015 *< β <* 0.0333 day^−1^ (depending on the value of the cost, *ϕ*_*C*_). Below the dotted line in the pink bistable region, there is bistability between the “all control” and DFE; above, there is bistability between the two-strategy equilibrium and the DFE.

Thus, as the proportion using resistance reaches increasingly high levels, fewer noncontrollers will switch to using the control scheme.

When the improved crop was tolerant, however, such an “all control” equilibrium was possible (Fig. 3b). The primary benefit of tolerant crop is that, when infection occurs, there is a lower loss of yield compared to unimproved (or resistant) crop. As there is no reduction in the viral titre, tolerant crops are just as likely to act as sources of infection as unimproved crops. Thus, the benefits of planting tolerant crop are experienced by the grower that plants tolerant crop, so there is no disincentive for other growers to also use it.

We then further investigated the initial conditions that could lead to bistability. Fig. 4a shows the effect of the rate of horizontal transmission (*β*) on the stability of equilibria. There is a bistable region between *β* = 0.02 and 0.0333 day ^−1^, and which equilibrium (either the disease-free or all-control equilibrium) is attained will depend on initial conditions. The discontinuous system leads to kinks in the graphs, whose ordering follows that outlined in Appendix 2.

**Figure 4:**
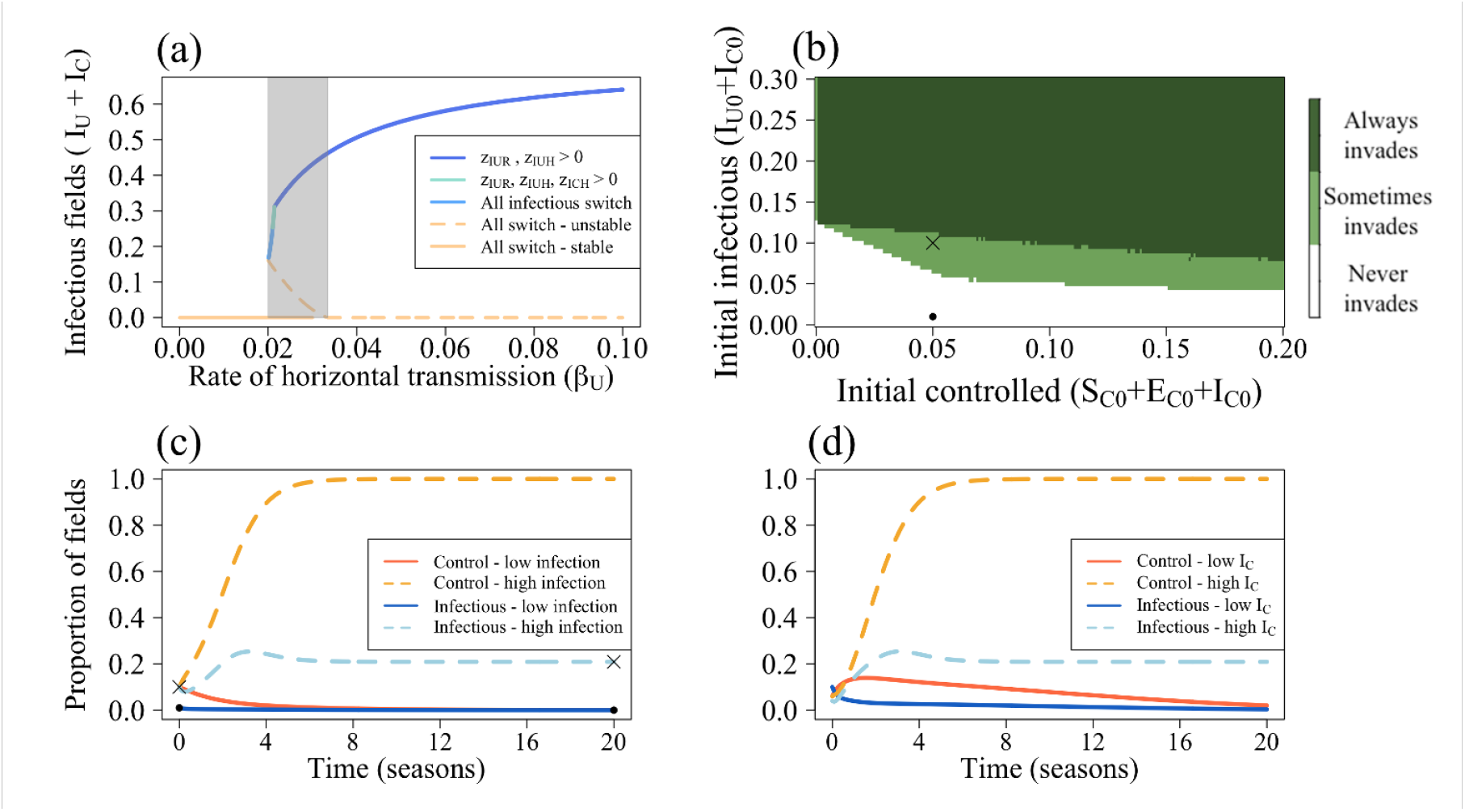
Bifurcation for the tolerant parameterisation. (a) Between *β* = 0.02 and 0.0333 day ^−1^ (the grey region), two biologically-meaningful equilibria are possible. Below *β* = 0.02 day ^−1^, only the disease-free equilibrium (DFE) is stable; above *β* = 0.0333 day ^−1^ (when *R*_0_ *>* 1), only the disease-endemic equilibrium is. The kinks in the graph are caused by the change in form of the switching terms. (b) The effect of the initial proportion of tolerant fields (*S*_*C*0_ + *E*_*C*0_ + *I*_*C*0_) and infectious fields (*I*_*U*0_ + *I*_*C*0_) on the persistence of disease at equilibrium. To only account for these factors, *E*_*U*0_ = *E*_*C*0_ = 0. When *S*_*C*0_ + *E*_*C*0_ + *I*_*C*0_ and *I*_*U*0_ + *I*_*C*0_ are very low, all initial conditions lead to the DFE. As these quantities increase, so too does the probability of disease persistence. The black cross and dot denote the proportions used in (c). (c) When *I*_*U*0_ + *I*_*C*0_ = 0.01 (“low infectious”), disease died out. However, once *I*_*U*0_ + *I*_*C*0_ = 0.1, disease persisted. (d) For the “high infection” parameterisation from (c), we investigated the effect of the proportion of infectious improved crop.When 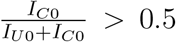 (as (*S*_*U*0_, *E*_*U*0_, *I*_*U*0_, *S*_*C*0_, *E*_*C*0_, *I*_*C*0_) = (0.83, 0, 0.03, 0.03, 0, 0.07); “high *I*_*C*_”), disease can persist. However, if 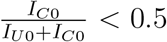 (as (*S*_*U*0_, *E*_*U*0_, *I*_*U*0_, *S*_*C*0_, *E*_*C*0_, *I*_*C*0_) = (0.83, 0, 0.07, 0.07, 0, 0.03); “low *I*_*C*_”), disease dies out. Other than *β* = *δ*_*β*_*β* = 0.0028 day^−1^ (which is within the bistable region), parameters are as in Table 2.

The initial proportions of growers using tolerant crop (*S*_*C*0_ + *E*_*C*0_ + *I*_*C*0_) and infectious fields (*I*_*U*_ + *I*_*C*_) had a large impact on the final proportion of infectious fields. We did this for *β* = *δ*_*β*_*β* = 0.0028 day^−1^, which is within the bistable region in Fig. 4a. At very low levels of both initially controlled and infectious fields, the system always goes to the disease-free equilibrium (DFE), with no disease persisting (Fig. 4b). As both increase, disease is more likely to invade until both controlled and infectious fields are sufficiently high that the system will always go to a diseaseendemic equilibrium.

We compared two sets of initial conditions, which differed in their initial proportion of infectious fields. In Fig. 4c, *S*_*C*0_ + *E*_*C*0_ + *I*_*C*0_ = 0.1 (i.e. initially 10% of fields are planted with improved crop). In the “high infection” scenario, 10% of fields are infectious; for the “low infection” scenario, this is just 1%. In the former scenario, disease persisted at equilibrium, whilst it died out in the latter.

The ratio of initially-infectious improved and unimproved crop (Fig. 4d) was important in determining disease persistence. If 70% of initially-infected fields were tolerant ((*S*_*U*0_, *E*_*U*0_, *I*_*U*0_, *S*_*C*0_, *E*_*C*0_, *I*_*C*0_) = (0.83, 0, 0.03, 0.03, 0, 0.07); “high *I*_*C*_”) disease persisted as it did in Fig. 4c. However, if there were fewer infectious tolerant fields ((*S*_*U*0_, *E*_*U*0_, *I*_*U*0_, *S*_*C*0_, *E*_*C*0_, *I*_*C*0_) = (0.83, 0, 0.07, 0.07, 0, 0.03); “low *I*_*C*_”), disease died out. Though this is not the only condition for disease persistence when *R*_0_ *<* 1, as at high levels of *I*_*U*0_ disease can persist even if there are no *I*_*C*0_ fields, these results cumulatively suggest that the presence of infected tolerant crop early in the epidemic can lead to alternative equilibrium being attained. This is possibly driven by the lower probability of detection of tolerant crop (*δ*_*v*_*v* = 0.1), which means that it is not removed quickly once it becomes infectious and allows disease to spread.

### 4.3 Q3: Impact of tolerant or resistant crop on grower behaviour

#### 4.3.1 Effect of epidemiological and economic parameters on grower behaviour

We first investigated the impact of changes to the rate of horizontal transmission on the adoption of improved crop. For all values of the rate of horizontal transmission in non-improved crop (*β*), the proportion of infected fields was higher when the improved crop was tolerant than when it was resistant (Fig. 5a *vs* b). Disease could also invade at a lower value of *β* (in the bistable region where *R*_0_ *<* 1). The expected profits for both strategies were higher when there was resistant crop (Fig. 5c,d), and the non-controllers gained more benefit from others using the resistant crop than from the tolerant crop.

**Figure 5:**
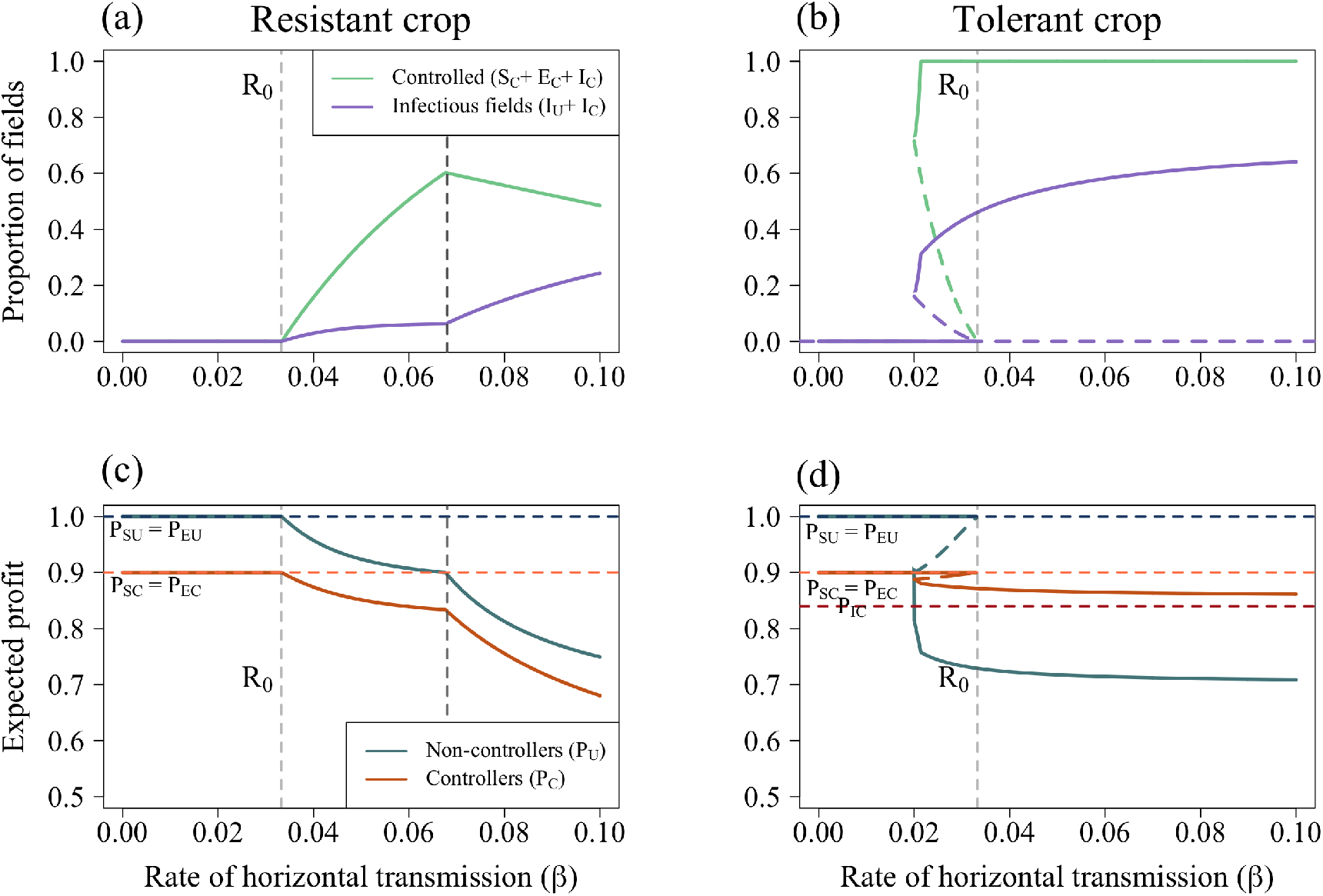
Response to the rate of horizontal transmission in non-improved crops (*β*). (a) Change in controllers and infectious fields when the improved crop has resistant characteristics. Generally, with an increase in *β* comes an increase in *C*, though at *β* = 0.067 day^−1^ there is a decrease and corresponding increase in infectious fields. (b) The change in proportion of growers using the improved crop (*C*) and infectious fields (*I*_*U*_ + *I*_*C*_) when the improved crop has tolerant characteristics. (c) Change in the expected profit for non-controllers (*P*_*U*_, Equation 35) and controllers (*P*_*C*_, Equation 36) for resistant crop. Until *β* = 0.067 day^−1^, controllers with susceptible or latently-infected crop should switch strategy as *P*_*U*_ *> P*_*SC,EC*_. After this point, they should stop switching. As *P*_*ICH,ICR*_ *< P*_*U*_, these growers should always change strategy, leading to a fall in the proportion of controllers. (d)Change in the expected profit for non-controllers (*P*_*U*_) and controllers (*P*_*C*_). For each value of *β*, growers managing susceptible or latently-infected fields of neither strategy should change (as *P*_*U*_ *< P*_*SC,EC*_ and *P*_*C*_ *< P*_*SU,EU*_). Other than those being varied parameters and initial conditions are as in Tables 2 and 3 respectively.

Generally, for both crop varieties, as *β* increased so too did the proportion of fields controlling. With the tolerant improved crop, this lead to a higher proportion of infectious fields (*I*_*U*_ + *I*_*C*_; Fig. 5b), as tolerant crops have a reduced probability of being rogued so there is a higher disease pressure on other fields in the system. Once disease invades at no point should growers of either strategy that have susceptible or latently-infected fields switch strategy as *P*_*U*_ *< P*_*SC,EC*_ and *P*_*C*_ *< P*_*SU,EU*_.

The trend is broadly similar for resistant crop. However, as *β* increases, there is a decrease in the proportion controlling. As *P*_*U*_ approaches *P*_*SC,EC*_, fewer growers managing fields of these types should switch strategy. However, the increase in infection pressure means that more of these fields will become infected, achieving the lowest payoff. Thus, the growers have a non-zero probability of switching strategy.

Once *β >* 0.067 day^−1^ (marked by the vertical dashed line in Fig. 5), the proportion of controllers falls. At this point, the high infection pressure means the expected profits of non-controllers falls below that of susceptible or latently-infected controllers, so they do not switch strategy (*P*_*U*_ *> P*_*SC*_ = *P*_*EC*_). Yet the fields of these controllers are still likely to get infected as resistance is incomplete, so the growers will incur the double penalty of the cost of control and loss due to disease. Thus, as *P*_*ICH,ICR*_ *< P*_*U*_ for all values of *β*, growers managing infected fields with improved crop should always consider switching strategy. As there are now fewer fields planted with resistant crop and the overall disease pressure increases, so too does the number of infectious fields.

The response to infection rate changed for different values of the cost of control, *ϕ*_*C*_ (Fig. 6). When the parameterisation was tolerant (Fig. 6a), bistability existed for values of *ϕ*_*C*_ *<*= 0.3, though the region where bistability was possible narrowed as the cost of control increased. When *ϕ*_*C*_ = 0.1 or 0.2, a scenario where grower behaviour meant that only tolerant crop was possible. Even at very high costs of control (*ϕ*_*C*_ = 0.5), some growers nevertheless controlled at equilibrium.

**Figure 6:**
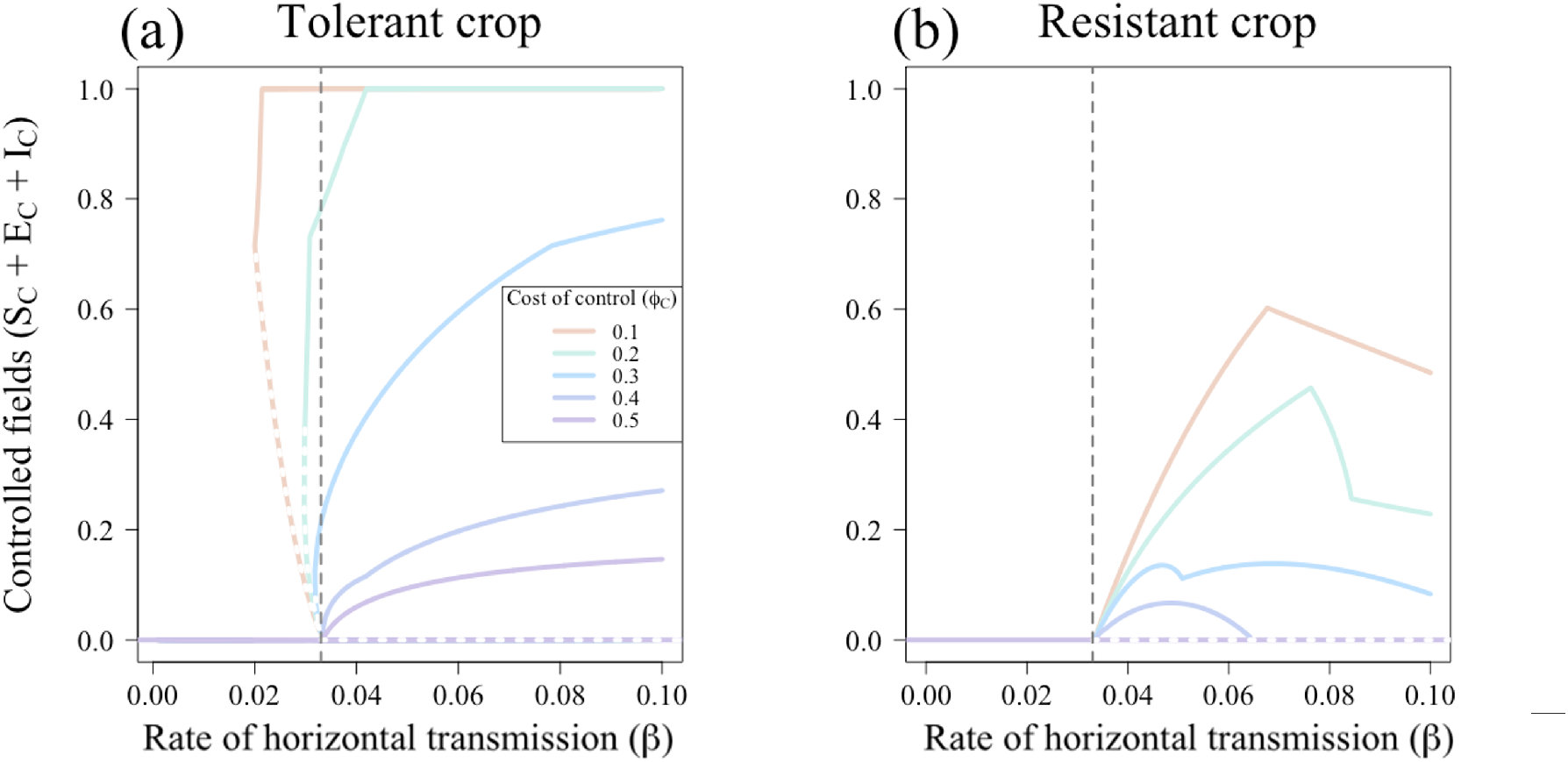
Effect of the cost of control (*ϕ*_*C*_) on uptake of improved crop. The grey dashed lines show where *R*_0_ = 1), and grey dots show unstable equilibria. (a) When improved crop is tolerant, when *ϕ*_*C*_ *≤* 0.3, bistability is possible. The range of *β* values for which bistability exists is larger with a lower *ϕ*_*C*_. Even at very high costs of control (*ϕ*_*C*_ *>* 0.4), control persists at equilibrium. (b) When improved crop is tolerant, no bistability exists. Only once *R*_0_ *>* 1 can control persist. However, as *β* gets larger, even at low costs of control fewer growers use resistant crop. Note, the maximum possible value for *ϕ*_*C*_ when the crop is resistant is *ϕ*_*C*_ = 0.4, otherwise payoff for infected resistant crop, *P*_*IUH*_, would be negative for the default parameterisation.

When the improved crop is resistant, disease can only invade once *R*_0_ *>* 1, irrespective of the cost of control. Once disease invades, control only persists for a narrow range of *β*, as above a certain threshold, control is seen as too costly (as growers are more likely to pay the dual penalty of the cost of control and loss due to disease, *L*_*C*_, thus earning the lowest possible profit, *P*_*ICH*_). The range for which resistant crop is used at all is narrower for larger values of *ϕ*_*C*_. The kinks in this graph (such as in Fig. 6a when *ϕ*_*C*_ = 0.3 and *β* = 0.78 day^−1^ or Fig. 6b when *ϕ*_*C*_ = 0.1 and *β* = 0.72 day^−1^) are caused by changes in the switching terms, discussed in detail in Appendix 6 Fig. 1.

#### 4.3.2 Comparison of profits for tolerant and resistant crop

We now compare the expected profits at equilibrium for controllers and non-controllers when the improved crop is either tolerant or resistant (Fig. 7, which shows the difference in expected profit for non-controllers (a) and controllers (b) when crop is resistant and when it is tolerant). The expected profits for unimproved, tolerant and resistant crop is shown in Appendix 6 Fig. 3. When generating these graphs, we chose initial conditions that would always guarantee a disease-endemic equilibrium in the bistable region to ensure that differences are calculated between equivalent equilibria (*I*_*U*0_ + *I*_*C*0_ = 0.15, *S*_*C*0_ + *E*_*C*0_ + *I*_*C*0_ = 0.2, Fig. 4b).

**Figure 7:**
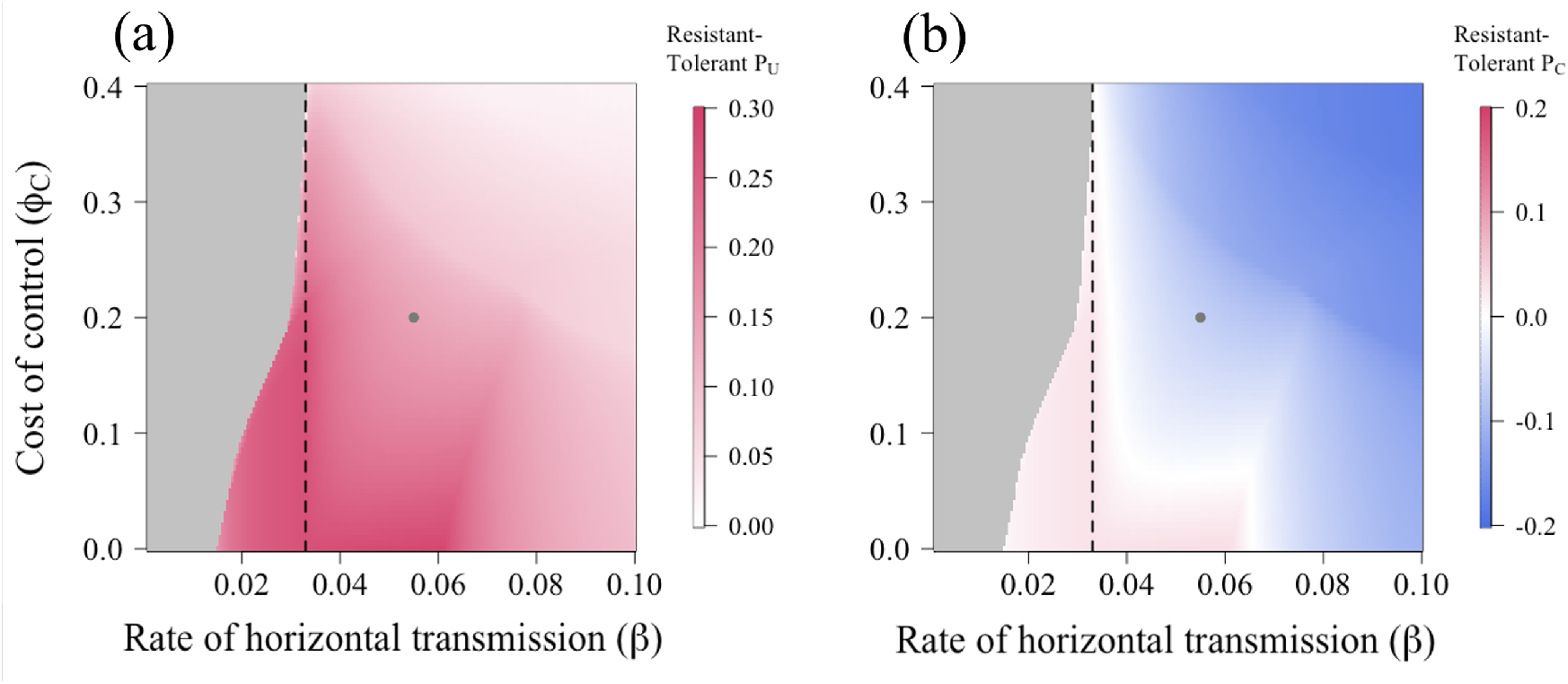
Difference in expected profits when there is resistant crop vs. tolerant crop. (a) The expected profit for unimproved crop and (b) the expected profit for improved crop. The grey dots indicate the default parameterisation and the white section below *R*_0_ = 1 (the black dashed line) is where there is no difference in profit due to the lack of improved crop being used. Other than those being varied, parameters and initial conditions are as in Tables 2 and 3 respectively.

Directly comparing the types of improved crop and their effect on the expected profits we can see that for non-controllers, the presence of the improved crop in the system is always more beneficial when crop is resistant (Fig. 7a). However, whether tolerant or resistant crop improves controllers’ profits depends on parameter values (Fig. 7b). At low-to-medium costs of control (*ϕ*_*C*_) and transmission rates (*β*), those growing resistant crop earn slightly more than they would if they used tolerant crop. Both improved crop types cost the same amount (*ϕ*_*C*_), so it is more advantageous to use resistant crop when there is already a low probability of infection, and effectively completely avoid incurring the yield loss due to infection, than to be tolerant and risk losing yield (even if the yield loss itself is small).

In both graphs, the increase and subsequent decrease in the benefit of resistance for non-controllers and resistant controllers that begins when *β* = 0.064 day ^−1^ in Figs. 7a,b is caused by changes in the switching terms (Appendix 2) and changes in infection pressure.

## 5 Discussion

Though tolerance and resistance to disease have been widely researched in plant biology, little thought has been given to the distinct epidemiological consequences of deploying tolerant or resistant varieties upon a community of growers, and none to how this affects the decisions of growers to partake in control. Growers’ behaviour more broadly, and its effects on epidemic outcomes remains largely overlooked in plant disease epidemiology (with some exceptions including Milne et al. (2016), McQuaid et al. (2017a), Szyniszewska et al. (2019), Milne et al. (2020), Bate et al. (2021), Saikai et al. (2021) and Murray-Watson et al. (2022)). Here we investigated the effect of a fixed proportion of “improved” crop (with either tolerant or resistant characteristics) on growers’ profits and subsequently how this affected growers’ use of improved crop when given the choice. Though the models we developed can in principle be applied to a broad range of pathosystems, we demonstrate the model using TYLCV as a case study.

As is intuitive from the underlying epidemiology, when a fixed proportion of growers was assigned tolerant crop, there was an increase in the proportion of infectious fields compared to when the improved crop was resistant (Fig. 2b *vs*. d). However, for the tolerant crop, the negative impact of having more infectious fields was predominantly felt by the growers using unimproved crop, who saw a larger fall in profits than the controllers. This was due to the negative externalities generated by the use of tolerant crop; as tolerant crop has reduced symptom development, it has a lower rate of removal via roguing. This allows more disease to build in the system, increasing the probability of infection for fields with unimproved crop and consequently reducing the profit of non-controllers.

When there was a constant proportion of growers using each strategy, when the proportion using resistant crop, *C >* 0.55, disease goes extinct (Fig. 2d). Despite the increase in profits experienced by all growers when disease was eliminated via the use of the resistant crop, most of the benefit was experienced by non-controllers, who saw a more substantial increase in profits than the controllers (Fig. 2b). This echoes past studies showing that if some proportion of growers in a landscape do control for disease, the benefits are widely felt (Hutchison et al. (2010),van den Bosch & Gilligan (2003), Lo Iacono et al. (2013)). In our case, growers of resistant crop generated positive externalities, benefiting others whilst incurring a cost themselves. Thus, the resistant crop is more beneficial to growers of both strategies than tolerant crop (which cannot be used to eliminate disease and increase the payoffs of non-controllers).

The nature of these contrasting externalities suggests that, given the choice, relatively few growers should choose to use resistant crop when available as they will gain more benefit when they do not (i.e. they will “free-ride” off of the costs incurred by controllers). Similarly, it suggests growers using tolerant crops incentivise others to do so too, as they will have higher yields when infected. To investigate these dynamics, we included growers’ behaviour in our disease spread model, with growers evaluating profitability based on the “grower vs. alternative strategy” method described in Murray-Watson et al. (2022) (which takes the same form as decision models in Milne et al. (2016), McQuaid et al. (2017a) and Saikai et al. (2021)). In this model, growers compared their own outcome from the previous season (i.e. whether they had a rogued field that had resistant crop, a susceptible field with unimproved crop *etc*.) with the average expected profit of the alternative strategy. The model is based on “strategic-adaptive” expectations, which balances a grower’s previous experience against the probability of future events (Fenichel & Wang (2013)).

Once growers’ behaviour is introduced, the threshold proportion of growers using resistant crop needed to eliminate disease is never reached, even at high disease pressures (Fig. 3a). This is because of “free-riding”; growers gain more benefit from the protection provided by others using resistant crop than they would if they themselves used resistant crop (in Fig. 5c, the expected profit for non-controllers is higher than for controllers (*P*_*U*_ *> P*_*C*_)). Thus, though the use of resistant crop can theoretically lead to disease elimination, when considering the behaviour of growers it is not possible.

When the improved crop was tolerant, however, bistability between disease-free and disease-endemic equilibria was observed when the basic reproduction number (*R*_0_) was less than unity (Fig. 3b). Previous epidemiological models including factors such as imperfect vaccination, risk-structure or re-infection (e.g. Arino et al. (2003), Gumel & Song (2008), summarised in Gumel (2012)), vector dynamics (Garba et al. (2008), Cunniffe et al. (2022)), fungicide application (Castle & Gilligan (2012)) or aspects of individual behaviour (Ajbar et al. (2021), Hadeler & Castillo-Chavez (1995)) have also identified such bistable regions. In our case, changing the rate of horizontal transmission (*β* = *δ*_*β*_*β* for the tolerant parameterisation) induced this bistability, and whether the system went to a disease-free or disease-endemic equilibrium depended on the initial proportion of infectious fields and initial proportion of controllers (Fig. 4b). Increasing the cost of control reduced the size of the region in parameter space in which bistability was observed (Fig. 6a). If the cost of control is sufficiently high such that tolerant crop is never more profitable than unimproved crop, the bistable region is eliminated. Bistability was not observed when the improved crop had the default resistant parameterisation (Figs. 3a and 6b). Thus, the use of tolerant crop may lead to less predictable outcomes at lower values of *δ*_*β*_*β*, as having a *R*_0_ *<* 1 is no longer sufficient to prevent disease spread.

When growers’ behaviour was introduced, an “all control” equilibrium, where all growers used improved crop, attained at low rates of horizontal transmission for the tolerant parameterisation (Fig. 5b). As seen for the fixed proportions, however, this was accompanied by a higher number of infectious fields and correspondingly low expected profit for non-controllers (Fig. 5a). It is these low expected profits for noncontrollers that drive the higher participation in control, as the profits for those using tolerant crop remain high irrespective of the infection pressure. The benefits of using tolerant crop are felt privately by those using it, generating negative externalities for others.

Conversely, such an “all control” equilibrium was not achieved when the improved crop was resistant, since it generates positive externalities for non-controllers (Figs. 3a and 5b). This is a product of both how the model was set up and the nature of the externalities produced. As the lowest possible payoff was achieved by growers of resistant crop that did not rogue their fields (*P*_*ICH*_), any of these growers should always have a non-zero probability of switching strategy (*z*_*ICH*_ *>* 0). Thus, there will always be non-controllers at equilibrium. Additionally, the reduced probability of infection meant that the need to control was reduced, disincentivising growers from switching to the costly control strategy. This conflict between private and social benefits is often observed in epi-economic models and is theorised to be the reason why many vaccination schemes fail to achieve a socially optimal level of vaccination (Brito et al. (1991), Geoffard & Philipson (1997), Sadique (2006)).

For growers who choose to use improved crop, tolerant varieties generally give better outcomes than resistant ones (except in cases where infection is unlikely enough that the probability of incurring the loss due to disease is low) (Fig. 7b). This will generate lower payoffs for non-controllers, who earn higher profits when there are more resistant fields (Fig. 7a). In our model, there was widespread use of control when the tolerant crop was effective at reducing yield loss and not too costly as to discourage control when the probability of infection is low. However, this came at the cost of increasing the level of infection in the system and reducing the profits for non-controllers.

Several simplifying assumptions were made during this investigation. Our model was deterministic and did not account for spatial effects, both of which can influence epidemic outcomes. In our behavioural model, we assumed that all growers would have access to the same information regarding disease pressure and the expected profits. In reality, a grower’s knowledge of these quantities will be highly dependent on their communication network (Milne et al. (2016)), their trust in expert knowledge (Sherman & Gent (2014)), their experience with previous outbreaks (Garcia-Figuera et al. (2021)) *etc*.. How growers react to differences in profit (represented by our parameter *η*) will also vary between individual growers, and will impact long-term outcomes (Murray-Watson et al. (2022)). Growers must balance these information sources with market demands to make their decisions regarding disease control, and become what Kaup terms the “reflexive producer” (Kaup (2008)).

Overall, this study has shown that tolerant and resistant varieties of crop have different effects on disease outcomes, and provide benefits to different groups of growers (controllers *vs*. non-controllers). In particular, even when resistant crop was available, disease was never eliminated from the system (even though it was theoretically possible) as too few growers chose to use the resistant variety. This is an important consideration as previous studies have found optimal cropping ratios for different sets of conditions (e.g. Ohtsuki & Sasaki (2006)); in reality, the strategic decision making of growers, as well as other factors such as their access to information or risk aversion, may mean that these ratios are never attained. Accounting for these behaviours can help improve future models of control uptake and in turn our understanding of how plant diseases spread.

## 6 Acknowledgements

REMW acknowledges the Biotechnology and Biological Sciences Research Council of the United Kingdom (BBSRC; https://bbsrc.ukri.org/) for support via a University of Cambridge DTP PhD studentship (Project Reference 2119272).

## 7 Author contributions

## 8 Data availability

## 10 Appendix 1

Parameterisation

The rate of horizontal transmission, *β*, is parameterised such that after 10 seasons, 60% of fields are infectious. This gives a value of *β* = 0.055 day^−1^. To account for the reduced susceptibility of resistant plants, we set the parameter *δ*_*β*_ = 0.5 as an illustrative example (so “resistant” plants still have some probability of being infected, as would be the case for quantitative disease resistance French et al. (2016)). Resistant plants are less likely to act as sources of inoculum for whitefly vectors than susceptible plants (Lapidot et al. (2002), Legarrea et al. (2015)). We set the reduced probability of infection from an infectious, resistant field (*δ*_*σ*_) as 0.5 as an illustrative example of this phenomenon.

The cropping period, *γ*, is 120 days, in line with tomato cultivation regimes (Holt et al. (1999*a*), Rocco & Morabito (2016)).

Complete crop losses due to TYLCV have been historically reported in many regions (e.g. in the Middle East (Czosnek & Laterrot (1997)) and the United States of America (Fonsah et al. (2018))), though certain management practices can alleviate such extreme events. Use of improved cultivars when TYLC is present can increase yield by up to 40% (Vijeth et al. (2018), Riley & Srinivasan (2019)). Riley & Srinivasan (2019) evaluated tomato yield for a combination of commonly-used control methods. When grown with silver mulch and cyantraniliprole insecticide, a susceptible variety *FL47* had a yield of 47 kg per plot, which we use as a proxy for maximum yield. Without either of those treatments, the yield was 18 kg per plot. The best resistant variety (*Security*) had a yield of 50 kg and 28 kg per plot respectively. However, under our definition, *Security* is better defined as a tolerant variety rather than a resistant one, as it did not completely restrict viral replication. Using these values we can estimate the yield loss to be *≈* 60% for susceptible and resistant cultivars and up to *≈* 45% in tolerant cultivars. As such, *L* = 0.6 and *δ*_*L*_*L ≤* 0.45, though both of these parameters can be scanned over to account for environmental and cultivar effects.

The latent period 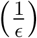, during which the host plant is infected but not infectious, can be estimated by using the number of days post infection when DNA can be detected. Significant amounts of viral DNA can be detected after 8 days (Ber et al. (1990)), which can be up to a week before symptoms appear. In Holt et al. (1999*b*), a latent period of 13 days was used. Here, we use an intermediate value of 1*/ϵ* = 10 days as the latent period of a single plant. Using the model stipulated in Holt et al. (1999*a*), though where the rate of harvesting and replanting is zero, if we then begin the season with 100% of plants in the *E* compartment, it takes around 41 days for 95% of these latently-infected plants to become infectious. We use this as our field-scale latent period (i.e. 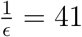 days).

We assume that symptoms and infectivity develop over roughly the same time scale (as symptoms can develop as early as two weeks after infection (Levy & Lapidot (2008)) and we use an individual-level latent period of 10 days). Thus we assume that the *P*_*SC*_ = *P*_*EC*_ and *P*_*SU*_ = *P*_*EU*_.

Economic analysis of tomato production in Georgia, USA estimates that the cost of improved cultivars is approximately 25% that of the total expected profits (Fonsah et al. (2018)), though these profit forecasts included other aspects of disease control. We set the value of the cost of control, *ϕ*_*C*_, as 0.1 of the total yield as an example of the extra costs control can entail, though our investigations involve a scan over possible costs.

The relative degree of tolerance and resistance will depend on both the unimproved and improved cultivars being compared. To allow for flexibility, the parameters presented in Table 2 are used to illustrate the effect of tolerance and resistance, though a range of parameters will be used in our investigations.

Similarly, the probability of symptom detection in an infectious field (*v*) will depend on a variety of anthropological, environmental and biological factors. The values presented in Table 2 are baseline parameters that can then be varied in our investigations. Importantly, when improved crop is tolerant, we assume that *δ*_*v*_*v < v*, whereas for resistant crop we assume *δ*_*v*_*v* = *v*.

For roguing to be worthwhile, it must reduce the potential losses to a grower. Though premature harvest of fruit can incur a yield penalty (between 16-19% for vine-ripened tomatoes, Davis & Gardner (1994)), we presume that if a grower notices a field is infectious and harvests it before the end of the growing season, it is overall more beneficial and their loss due to disease is reduced by some factor, *ϕ*_*R*_ *<* 1. The value of *ϕ*_*R*_ will vary with crop cultivar and environmental conditions; the value presented in Table 2 is illustrative though can be varied.

## 11 Appendix 2

Ordering of switching terms

The values of the switching terms are determined by the values of the profits laid out in Equations 9-16. The ordering will depend on the relative values of three key parameters: the loss due to disease for unimproved crop (*L*_*U*_), the loss due to disease for the improved crop (*L*_*C*_) and the cost of control (*ϕ*_*C*_).

If we assume that *L*_*U*_ *> L*_*C*_ and *L*_*U*_ *>* (*L*_*C*_ + *ϕ*_*C*_) (as for the tolerant parameterisation), the ordering of the profits is as follows:

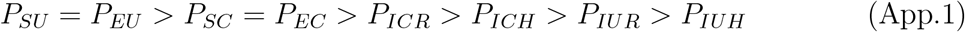

There are five possible combinations of values for the switching terms:

- *z*_*SC*_, *z*_*EC*_, *z*_*ICR*_, *z*_*ICH*_, *z*_*IUR*_, *z*_*IUH*_ *>* 0
- *z*_*ICR*_, *z*_*ICH*_, *z*_*IUR*_, *z*_*IUH*_ *>* 0
- *z*_*ICH*_, *z*_*IUR*_, *z*_*IUH*_ *>* 0
- *z*_*IUR*_, *z*_*IUH*_ *>* 0
- *z*_*IUH*_ *>* 0

If, instead, *L*_*U*_ = *L*_*C*_ and *L*_*U*_ *<* (*L*_*C*_ + *ϕ*_*C*_) (as for the resistant parameterisation and cases of the tolerant parameterisation with high values of *ϕ*_*C*_), the profits are ordered as:

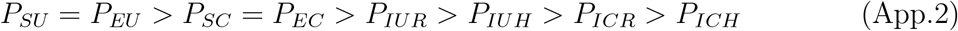

The following combinations of switching terms are possible:

- *z*_*SC*_, *z*_*EC*_, *z*_*IUH*_, *z*_*ICR*_, *z*_*ICH*_, *z*_*IUR*_ *>* 0
- *z*_*IUH*_, *z*_*ICR*_, *z*_*ICH*_ *>* 0
- *z*_*ICR*_, *z*_*ICH*_ *>* 0 *
- *z*_*IUR*_, *z*_*ICH*_ *>* 0 *
- *z*_*ICH*_ *>* 0

* The ordering of these terms depends on the proportional reduction in the loss of yield due to roguing (*ϕ*_*R*_).

## 12 Appendix 3

Mathematical details of non-behavioural model

We use the NGM method (van den Driessche (2017)) to calculate the basic reproduction number when there are two types of crop (improved and unimproved) present at the disease-free equilibrium, but growers cannot change strategy.

We focus only on the infected compartments, which are given by:

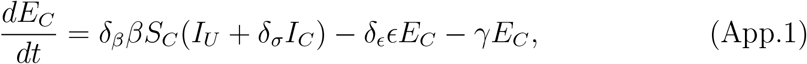

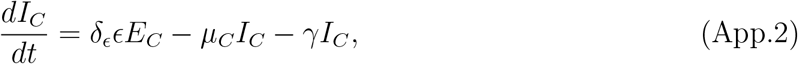

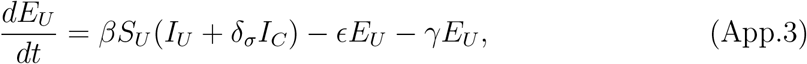

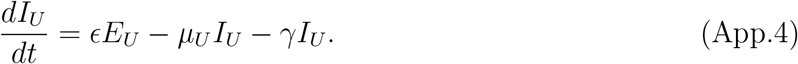

We first linearise these equations to give the Jacobian matrix and evaluate it around the disease-free equilibrium. We then decompose this Jacobian matrix into two further matrices: *F*, which is the matrix of terms relating to disease transmission, and *V* = *−Q*, where *Q* is the matrix containing non-epidemiological transition terms. The NGM, *K*, is then given by *FV* ^−1^ (van den Driessche (2017)).

For this system,

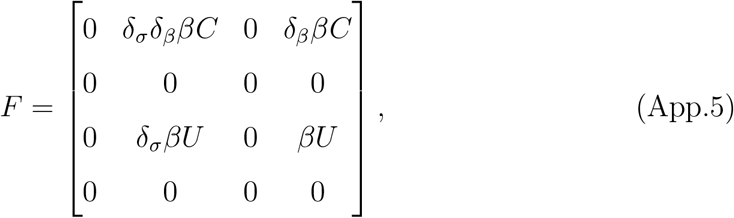

and

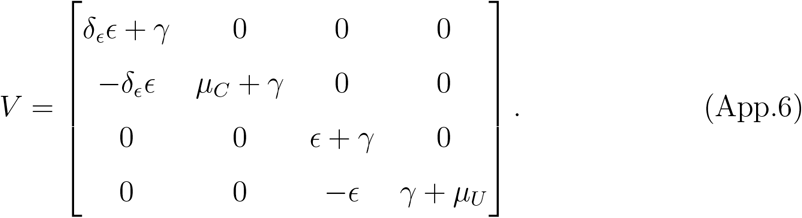

The inverse of *V* is given by:

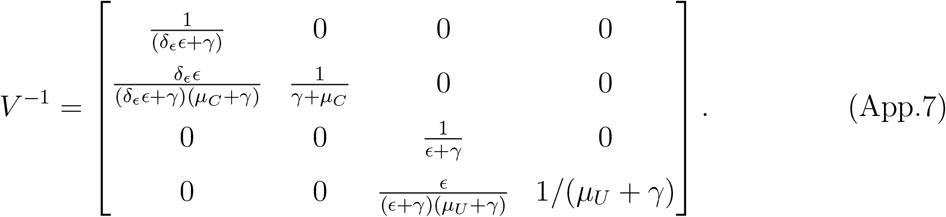

The NGM, *K* = *FV* ^−1^, is then given by:

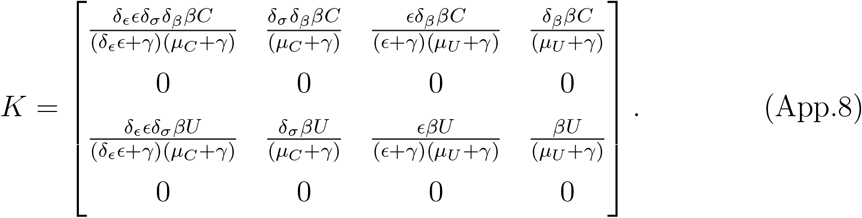

*R*_0_ is given by the leading eigenvalue of this matrix:

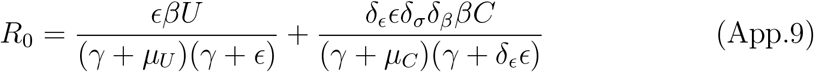

## 13 Appendix 4

Supplementary results for non-behavioural model

### 13.1 Parameters relating to tolerance and resistance

A broad range of parameter values relating to tolerant and resistant traits are possible depending on the cultivar and environmental conditions. As illustrative examples of the effect of changing parameters along the tolerance/resistance continuum, we investigate the effects of changing the probability of detection for improved crop (*δ*_*v*_*v*; Fig. 1) and relative susceptibility of improved crop (*δ*_*β*_; Fig. 2). In each case, profits were highest for both controllers and non-controllers when the parameterisation approached that of the resistant crop (i.e. a high probability of detection and low relative susceptibility). However, there was little impact on the profits of controllers who grew tolerant crop (Fig. 1c and Fig. 2c), as the low loss due to disease for tolerant crop means that the reduced probability of infection conferred by high *δ*_*v*_ and low *δ*_*β*_ is of little benefit.

Disease elimination was possible under many parameterisations for the resistant crop (due to the lower background infectivity of resistant crop, as well as its lower susceptibility and higher probability of infection). For the tolerant crop, when the relative susceptibility was very low, only then was disease elimination possible (Fig. 2a,c). This allowed growers to earn the maximum possible profits.

**Figure 1:**
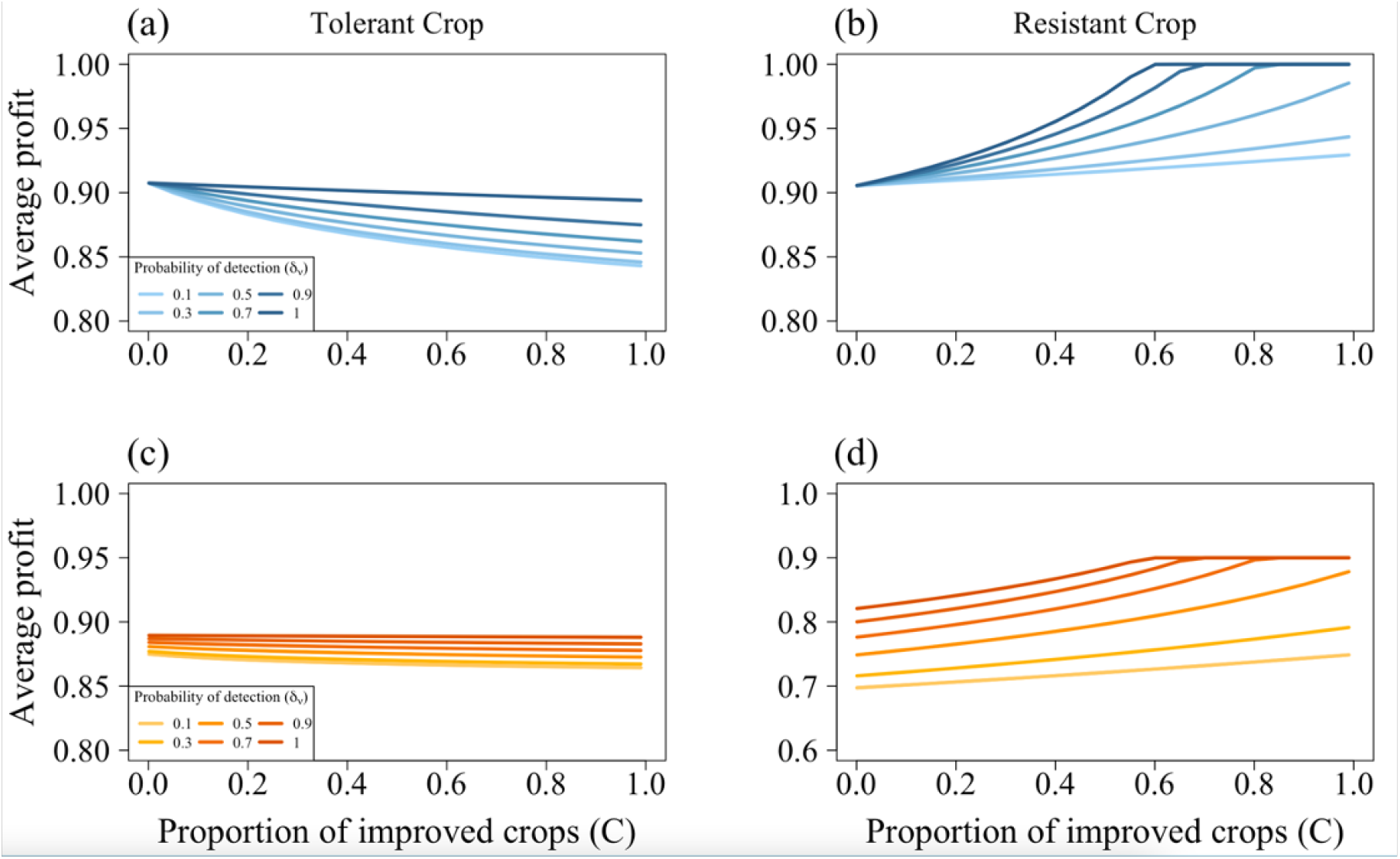
Change in average profits for tolerant and resistant parameterisation when the probability of detection for improved crop (*δ*_*v*_*v*) is varied. (a) and (c) show the average profit for unimproved and improved crop respectively for the tolerant parameterisation, whilst (b) and (d) show the same for the resistant parameterisation. In all cases, the highest profits were achieved when *δ*_*v*_ = 1 (i.e. infectious crop is always detected). The probability of detection had little impact on the average profit of controllers (c). This is because the probability of detection affects the rate at which infectious plants are removed; the higher the probability of detection, the lower the disease pressure and thus the lower the probability of incurring the loss due to disease. As the loss due to disease is low for tolerant crop, there is little impact on the profits of controllers. Additionally, disease was not eliminated when the crop was tolerant. When the crop was resistant, this also allowed disease elimination to occur at lower proportions of resistant crop (*C* = 0.6 when *δ*_*v*_*v* = 1, and (d)). Other than those scanned over, parameters are as in Table 2.

**Figure 2:**
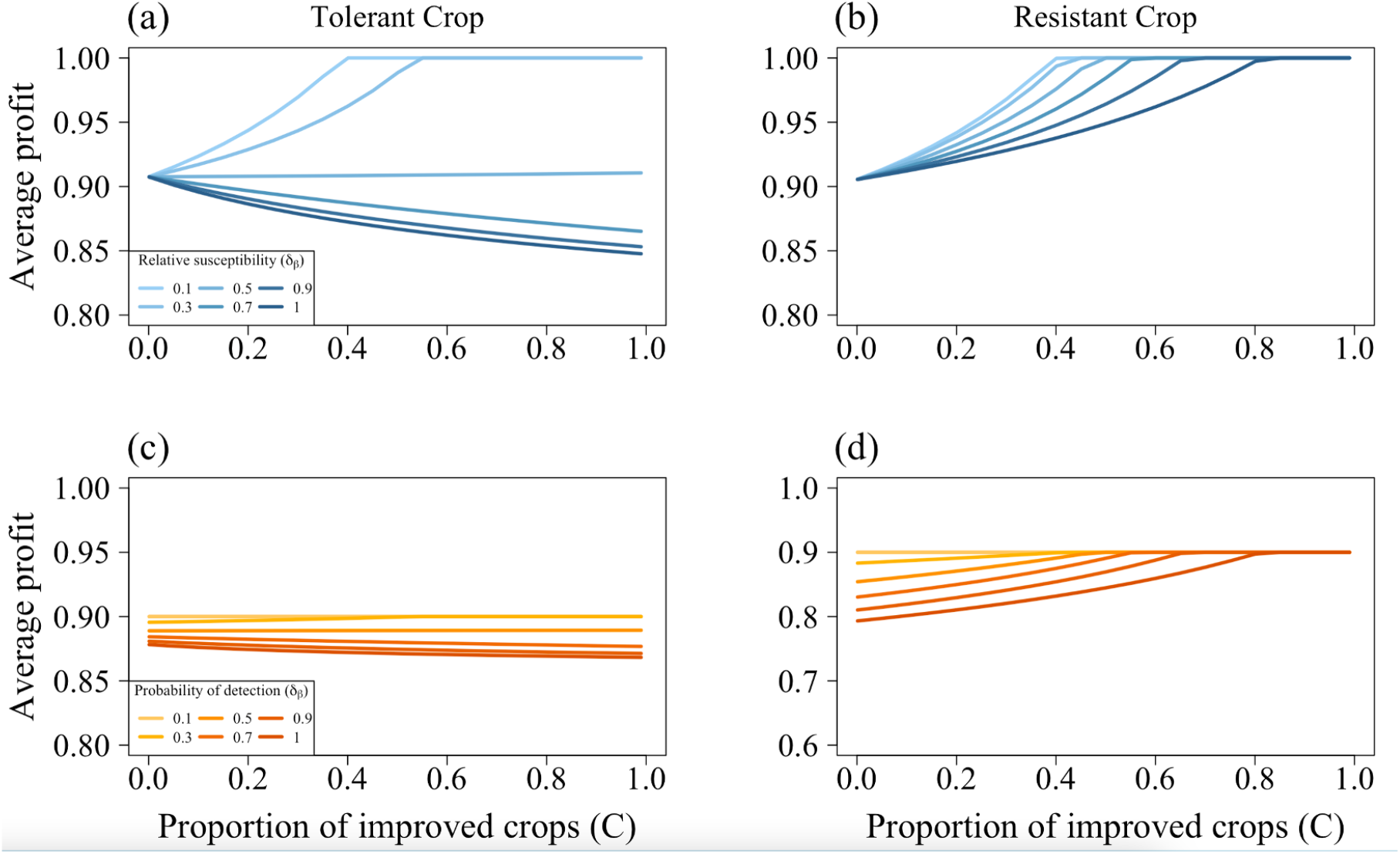
Change in average profits for tolerant and resistant parameterisation when the relative susceptibility of improved crop (*δ*_*β*_) is varied. (a) and (c) show the average profit for unimproved and improved crop respectively for the tolerant parameterisation, whilst (b) and (d) show the same for the resistant parameterisation. In all cases, at lower susceptibilities (which means the improved crop is less likely to become infected), profit increases. Indeed, under this parameterisation, disease can go extinct under the tolerant parameterisation (at *C* = 0.4 when *δ*_*β*_ = 0.1, (a) and (c)). There is little impact on the average profit for controllers when the improved crop is tolerant (c), as the low loss due to disease in the tolerant parameterisation means that the reduced probability of infection brought about by a lower *δ*_*β*_ have little effect. Disease elimination can occur in under all values of *δ*_*β*_ when the improved crop is resistant ((b) and (d)). Even at high values of *δ*_*β*_, the resistant crop still has a lower relative infectivity (*δ*_*σ*_) and higher probability of detection (*δ*_*v*_) than tolerant crop. Other than those scanned over, parameters are as in Table 2.

## 14 Appendix 5

Mathematical details of behavioural model

### 14.1 Evaluating stability for behaviour model

To investigate how the initial conditions affected the equilibrium, we conducted a randomisation scan with 10,000 sets of initial conditions and used *nleqslv* (Hasselman & Hasselman (2018)) in *R* to investigate the number of equilibria attained for each parameter set. The nature of the switching terms means that the system is discontinuous, and the equations will have a different form depending on the values of the state variables. There are ten possible Jacobians, depending on the values of the switching terms (Appendix 2). For each set of equilibrium values found from our randomisation scan, we evaluated the stability of that equilibrium using the appropriate Jacobian matrix.

### 14.2 Basic reproductive number of behavioural model

As the form of the equations differs between the model with fixed proportions and the behavioural model, *R*_0_ must be calculated separately for the behavioural model. For the disease-free equilibrium given by (*S*_*U*_, *E*_*U*_, *I*_*U*_, *S*_*C*_, *E*_*C*_, *I*_*C*_) = (*U*, 0, 0, 0, 0, 0), all switching terms should be non-zero. As there is no disease, there is no need for control, so no growers should use the control strategy. Additionally, any growers whose fields do become infected will have a lower payoff than the expected payoff of the alternative strategy (as there is no disease), so all non-controllers with infectious fields should switch strategy. The system of equations is therefore:

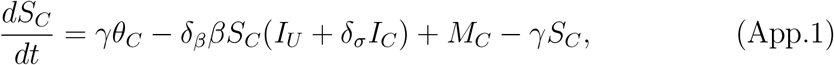

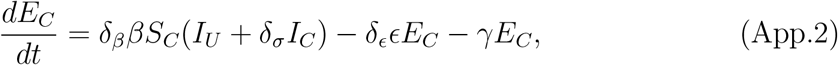

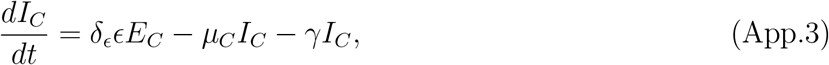

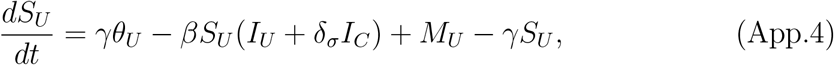

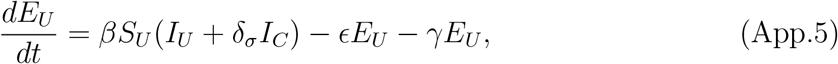

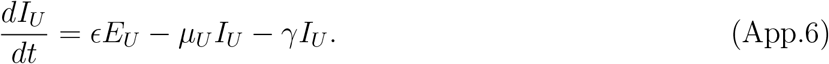

where:

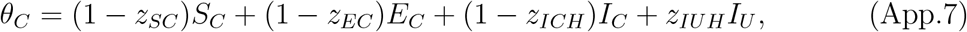

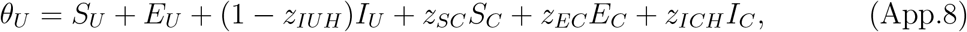

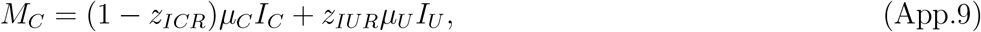

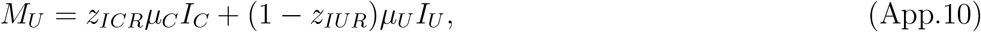

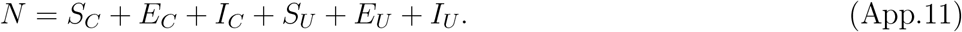

Using Equations App.1 - App.6, and the method outlined in van den Driessche (2017) and the main text, we find:

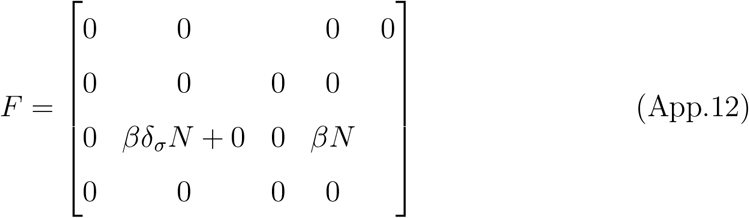

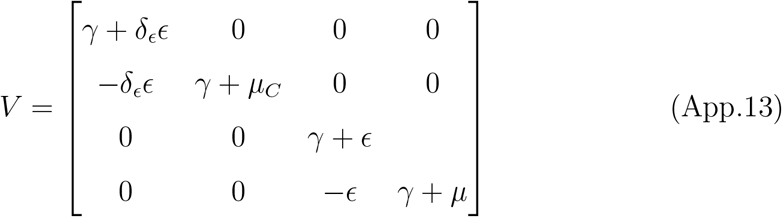

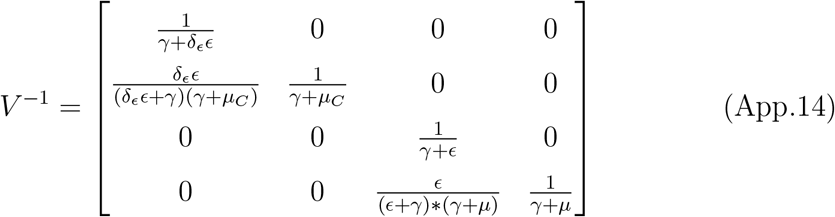

The NGM, *FV* ^−1^, can then be simplified to:

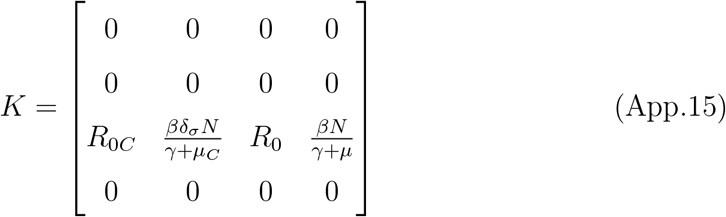

where 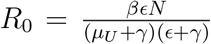 (Equation 58) and 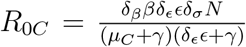 (Equation 59). The eigenvalue of *K* is given by *R*_0_, so the basic reproduction number is the same regardless of the whether behaviour is included in the model, depending only on whether *N* = *U* or *N* = *C*.

## 15 Appendix 6

Supplementary results for behavioural model

### 15.1 Underlying behaviour of switching terms for resistant parameterisation with variable cost of control

The kinks in Fig. 6b in the main text are caused by changes in the values of the switching terms. As the epidemic progresses, the expected profits for each strategy changes. If they fall below the profit for a particular outcome (for example, the profit for a controller with an infected field, *P*_*IUH*_), growers who have earned that outcome switch from having a non-zero probability of switching strategy to never switching. For different values of *ϕ*_*C*_, this occurs at different values of *β* (Fig. 1).

**Figure 1:**
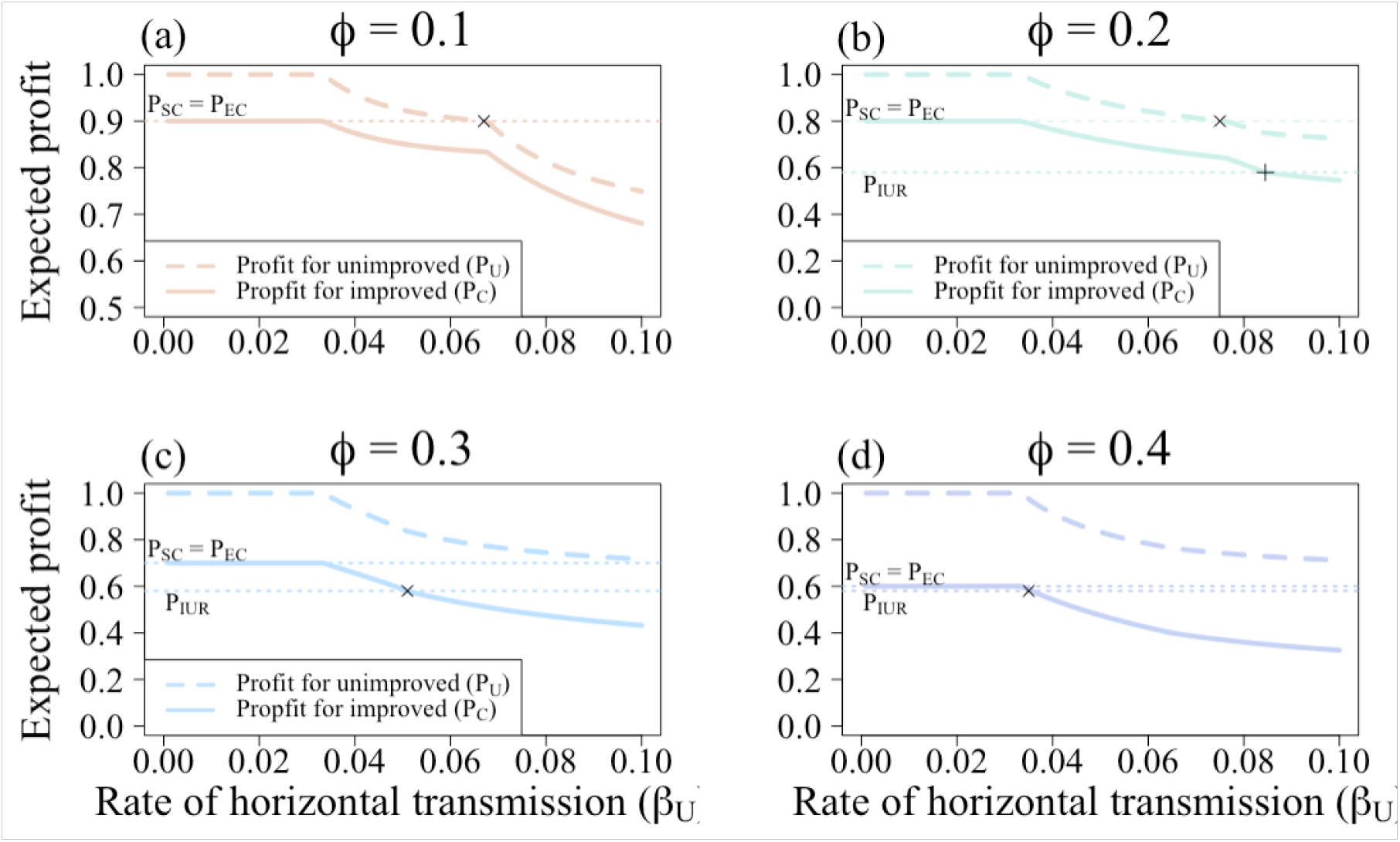
Change in expected profits with different values of the cost of control (*ϕ*_*C*_) for resistant parameterisation. (a) When *ϕ*_*C*_ = 0.1, the expected profits for non-controllers falls below the expected profits for controllers with susceptible or latently-infected (*S*_*C*_ or *E*_*C*_) crop at *β* = 0.068 day^−1^ (“x”). At this point, those with *S*_*C*_ or *E*_*C*_ fields stop switching strategy. (b) When *ϕ*_*C*_ = 0.2, *S*_*C*_ and *E*_*C*_ growers stop switching strategy at *β* = 0.075 day^−1^ (“x”). At *β* = 0.0845 day^−1^ (“+”), non-controllers with infectious fields that have been rogued also stop switching strategy (as *P*_*C*_ *< P*_*IUR*_). (c) When *ϕ*_*C*_ = 0.3, non-controllers with *I*_*UR*_ fields stop switching strategy at *β* = 0.051 day^−1^ (“x”), and controllers with *S*_*C*_ or *E*_*C*_ fields stop when *β* = 0.1 day^−1^. (d) When *ϕ*_*C*_ = 0.4, growers with *P*_*IUR*_ fields stop switching strategy at *β* = 0.035 day^−1^ (“x”). Controllers always have a non-zero probability of switching strategy for this parameter set. Other than those scanned over, parameters are as in Table 2.

### 15.2 Expected profits for unimproved, tolerant and resistant crop

The pattern outline below is the same as that for Fig. 5 in the main text, though for a two-way scan of the rate of horizontal transmission (*β*) and the cost of control (*ϕ*_*C*_).

Irrespective of whether the improved crop was tolerant or resistant, as *β* increased there was a corresponding increase in the proportion of infectious fields (*I*_*U*_ + *I*_*C*_; Fig. 2a,b). This increase occurred more quickly when the improved crop was tolerant, as tolerant crop has the same susceptibility and infectivity as unimproved crop, but is less likely to be detected and removed once infected.

Resistance is incomplete (Table 1 in the main text), so fields planted with resistant crop may still be infected. However, the reduced probability of infection means that there are overall lower proportions of infected fields. Participation in control is relatively high for low values of *β* and *ϕ*_*C*_, though this decreases as *β* gets larger (approaches 0.067 day^−1^ in Fig. 5 in the main text; where it occurs in Fig. 2a,c depends on the value of *ϕ*_*C*_). For these parameter values, *P*_*U*_ approaches *P*_*SC,EC*_ (Fig. 5a in the main text), so fewer controllers with susceptible or latently-infected fields should switch strategy. There is still a high infectious pressure, though, so more of these controlled fields will become infected. They will therefore incur the loss due to disease (*L*_*C*_), resulting in the lowest possible payoff. Expected profits for controllers (and consequently proportion of growers using control) briefly increases when *P*_*C*_ *< P*_*IUR*_ (which occurs at at *β* = 0.067 day^−1^ in Fig. 5a in the main text). The non-controllers who rogued their fields (achieving *P*_*IUR*_), should no longer switch strategy, and they replant *S*_*U*_ fields. There is still a high probability of infection, however, and many of these non-controlled fields will be infected by the time they are harvested. Some will be rogued before harvesting, preventing some loss of yield. Those that have not rogued achieve a low payoff and switch into the control strategy.

After this point, however, the proportion of controllers falls. The increased infection pressure means that the expected profit of non-controllers is lower than the profit of controllers with susceptible or latently-infected fields (*P*_*U*_ *< P*_*SC,EC*_). Controllers that harvest susceptible or latently-infected fields should therefore never consider switching strategy. The high infection pressure, however, means that many of these resistant fields will be infected before they are harvested. As *P*_*ICH,ICR*_ *< P*_*U*_ for these values of *β*, controllers with infectious fields should always switch strategy. As fewer growers control and plant resistant crop, the disease pressure increases and there are more infectious fields.

When the improved crop was tolerant, we chose initial conditions such that there would always be a disease-endemic equilibrium in the bistable region (*I*_*U*0_ + *I*_*C*0_ = 0.15, *S*_*C*0_ + *E*_*C*0_ + *I*_*C*0_ = 0.2, Fig. 4 in the main text). A high proportion of infectious fields was seen for most parameter combinations, in part due to the lower probability of infectious tolerant fields being removed by roguing. This accompanied a high degree of participation in control, as the low default value of *L*_*C*_ (= 0.06) and lower probability of paying the roguing cost. Additionally, once *R*_0_ *>* 1, costs of control *<* 0.2 “all-control” equilibrium, where *S*_*C*_ + *E*_*C*_ + *I*_*C*_ = *N*.

This “all-control” equilibrium was not seen in the parameterisation where the improved crop was resistant. This is both due to the positive externalities generated by the crop reducing the probability of infection for non-controllers and the structure of the model, which means growers with infectious resistant crop should always have a non-zero probability of switching strategy (as *P*_*ICR*_ us the lowest payoff).

**Figure 2:**
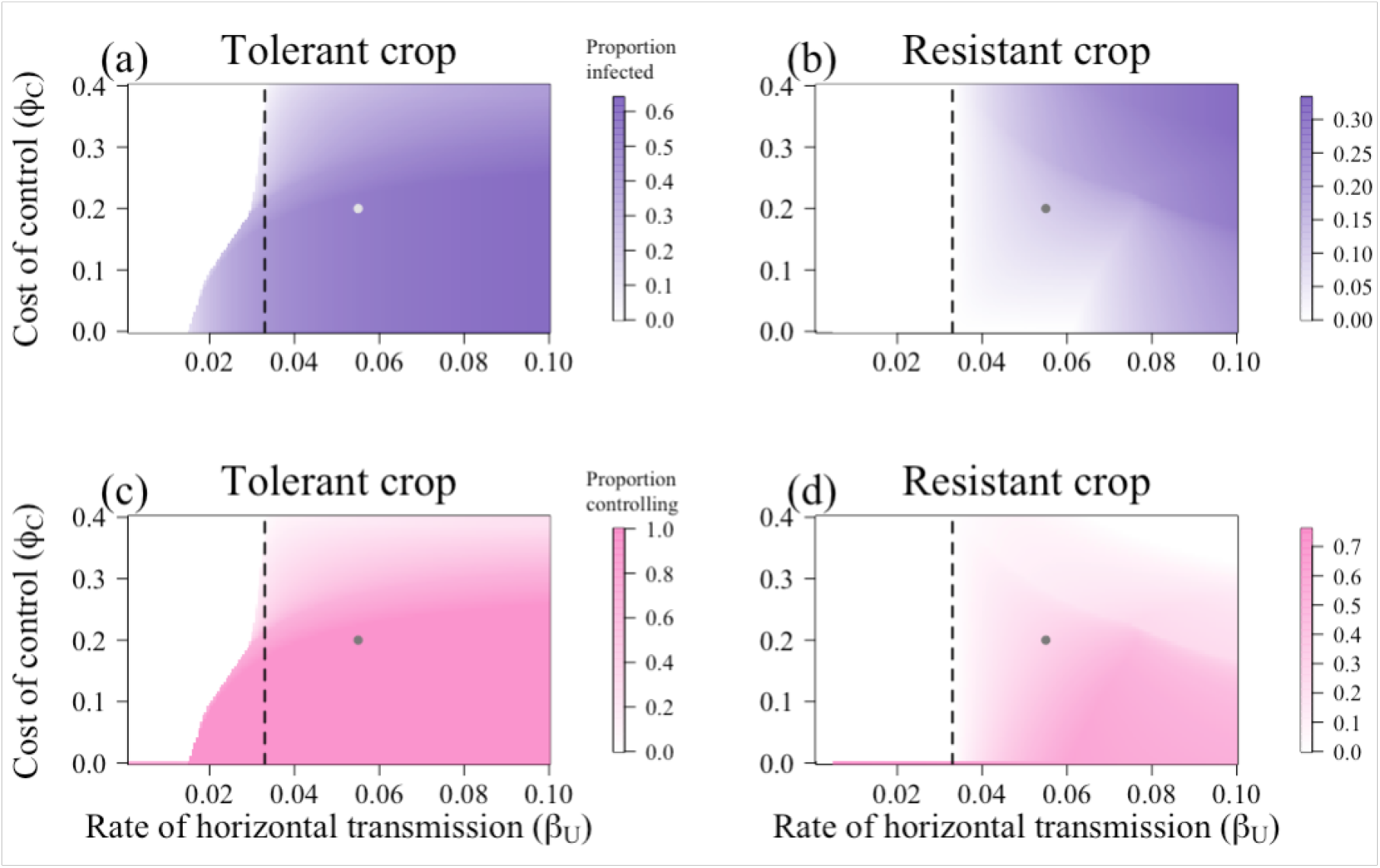
Response of the number of infectious fields and participation in control to the rate of horizontal transmission in non-improved crops (*β*) and the cost of control (*ϕ*_*C*_). (a) The change in proportion infectious fields (*I*_*U*_ + *I*_*C*_) and (b) change in participation in control when the improved crop has tolerant characteristics. Equivalent plots for resistant improved crop are shown in (c) and (d). In all graphs, the vertical dashed line at *β* = 0.0333 day ^−1^ is where *R*_0_ = 1. When the crop used by controllers tolerant to infection, there are high levels of infection and participation in control. Additionally, at low values of *ϕ*_*C*_, disease can invade when *R*_0_ *<* 1 (a). For resistant crop, there is a much lower proportion of infectious fields and controllers for most values of *β*. Other than those being varied, parameters and initial conditions are as in Tables 2 and 3 respectively.

The expected profits of both controllers and non-controllers follow a similar pattern to that of the proportion of infectious fields and controllers (Fig. 5 in the main text). In both cases, when *R*_0_ *<* 1 the profit of non-controllers (*P*_*U*_) is equal to that of susceptible/latently-infected non-controllers (except where bistability exists in the tolerant parameterisation). Once disease invades, profits of both controllers and non-controllers fall. However, for the majority of values of *β* and *ϕ*_*C*_, *P*_*U*_ is higher when there is resistant crop than when there is tolerant crop, indicating that non-controllers benefit more from the presence of resistant crop. However, the profits of growers using tolerant crops were generally higher than those using resistant crops, as the benefits generated by tolerant crops were experienced privately by the growers using them.

**Figure 3:**
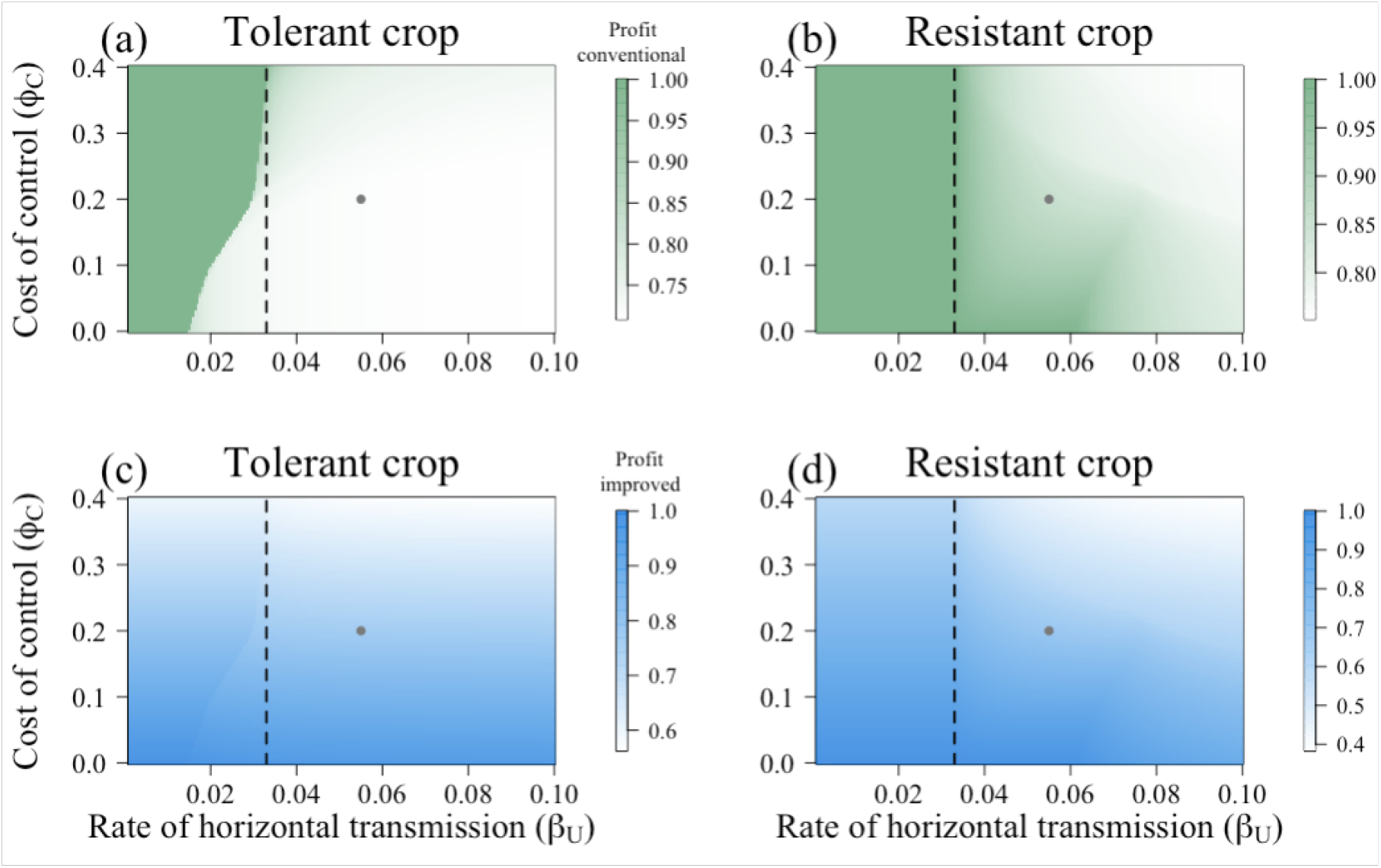
Response of the expected profits to the rate of horizontal transmission in non-improved crops (*β*) and the cost of control (*ϕ*_*C*_). (a) and (c) are the expected profits for non-controllers (*P*_*U*_) and controllers (*P*_*C*_) respectively when the improved crop is tolerant; (b) and (d) show the same for resistant crop. The highest profits for non-controllers are seen when *R*_0_ *<* 1, though *P*_*U*_ is generally higher for those in the resistant-crop scenario than the tolerant crop. Conversely, controllers that used tolerant crop generally had higher profits than those using resistant crop. The grey dots indicate the default parameterisation. Other than those being varied, parameters and initial conditions are as in Tables 2 and 3 respectively.

## Notes

### Competing Interest Statement

The authors have declared no competing interest.

